# Evolution of the *D. melanogaster* chromatin landscape and its associated proteins

**DOI:** 10.1101/284828

**Authors:** Elise Parey, Anton Crombach

**Affiliations:** Center for Interdisciplinary Research in Biology (CIRB), Collège de France, CNRS, INSERM, PSL Université Paris, 75005 Paris, France; (current address) Institut de Biologie de l’Ecole Normale Supérieure (IBENS), Ecole Normale Supérieure, CNRS, INSERM, PSL Université Paris, 75005 Paris, France; (current address) Inria, Antenne Lyon La Doua, 69603 Villeurbanne, France; Université de Lyon, INSA-Lyon, LIRIS, UMR 5205, 69621 Villeurbanne, France

**Keywords:** phylogenomics, chromatin-associated proteins, chromatin types, intron/exon structure, centromere drive, *D. melanogaster*

## Abstract

In the nucleus of eukaryotic cells, genomic DNA associates with numerous protein complexes and RNAs, forming the chromatin landscape. Through a genome-wide study of chromatin-associated proteins in *Drosophila* cells, five major chromatin types were identified as a refinement of the traditional binary division into hetero-and euchromatin. These five types were given colour names in reference to the Greek word *chroma.* They are defined by distinct but overlapping combinations of proteins and differ in biological and biochemical properties, including transcriptional activity, replication timing, and histone modifications. In this work, we assess the evolutionary relationships of chromatin-associated proteins and present an integrated view of the evolution and conservation of the fruit fly *D. melanogaster* chromatin landscape. We combine homology prediction across a wide range of species with gene age inference methods to determine the origin of each chromatin-associated protein. This provides insight into the evolution of the different chromatin types. Our results indicate that for the euchromatic types, YELLOW and RED, young associated proteins are more specialized than old ones. And for genes found in either chromatin type, intron/exon structure is lineage-specific. Next, we provide evidence that a subset of GREEN-associated proteins is involved in a centromere drive in *D. melanogaster*. Our results on BLUE chromatin support the hypothesis that the emergence of Polycomb Group proteins is linked to eukaryotic multicellularity. In light of these results, we discuss how the regulatory complexification of chromatin links to the origins of eukaryotic multicellularity.

## Introduction

The chromatin landscape consists of DNA, histones, and other associated proteins and RNAs, and plays a fundamental role in development, cellular memory, and integration of external signals. As a unique feature of the eukaryotic cell, it is closely tied to the evolution of eukaryotes, both regarding their origin and the major transition(s) to multicellularity (Newman 2005; Aravind et al. 2014; Gombar et al. 2014; Penny et al. 2014; Miyamoto et al. 2015; Sebé-Pedrós et al. 2017). At a basic level, chromatin is responsible for maintenance, organization, and correct use of the genome. Histone proteins package and condense DNA in the nucleus, and form a backbone for the action of structural and regulatory proteins. A variety of reversible post-translational modifications of histones, known as epigenetic marks, promote the recruitment of specific proteins. This creates a local context for nuclear processes such as transcriptional activity, replication, as well as DNA-repair. These and other epigenetic mechanisms involved in chromatin modification have been extensively characterized in a variety of eukaryotic species, which led to the observation that the chromatin landscape is effectively subdivided into a small set of distinct chromatin states (Filion et al. 2010; Ernst et al. 2011; Roudier et al. 2011). A largely open question, however, is how these chromatin states have evolved. In this work, we assess the evolutionary relationships of chromatin-associated proteins (CAPs) and present an integrated view of the evolution and conservation of the fruit fly *D. melanogaster* chromatin landscape.

Classically, chromatin is divided into two states, namely heterochromatin and euchromatin, the former a compacted DNA state in which transcription is mostly repressed and the latter an open, transcriptionally active configuration. This classification has been refined into multiple types of chromatin. In particular, a breakthrough result was presented by Filion et al., who established five major chromatin types in *D. melanogaster*, named with the colors YELLOW, RED, GREEN, BLUE, and BLACK. To do so, they used genome-wide binding profiles of CAPs obtained via DamID (Vogel, Peric-Hupkes, et al. 2007; Filion et al. 2010; van Bemmel et al. 2013). This approach is complementary to more commonly used genome-wide histone mark profiling techniques, such as ChiP-seq. Nevertheless, both are consistent with each other and serve as independent validation. Indeed, the five types can be mapped to an alternative classification into nine chromatin states, that is derived from histone modifications (Kharchenko et al. 2011).

The five chromatin types have different biological and biochemical properties. YELLOW and RED are two types of euchromatin. Looking at the CAPs that bind nearby transcription start sites (TSS), YELLOW mainly marks ubiquitously expressed housekeeping genes. In contrast, the genes with their TSS harbored in RED show more restricted expression patterns and are linked to specific tissues and developmental processes. Both euchromatin types are replicated in early S phase, and of the two, RED tends to be replicated first (Filion et al. 2010). GREEN and BLUE are two types of heterochromatin. GREEN is considered constitutive heterochromatin. It is identified by HP1-related proteins and is especially prevalent in pericentric regions as well as on chromosome 4. BLUE is facultative heterochromatin and concerns mostly genes specifically repressed during development. It is notably composed of the Polycomb Group (PcG) proteins, which were originally discovered in *D. melanogaster* to repress Hox genes and were later found to have a general role in development. (Lewis 1978; Duncan 1982; Boyer et al. 2006; Lee et al. 2006; Nègre et al. 2006). Finally, BLACK was originally interpreted as a major repressive chromatin type, but recent findings indicate it is better described as a near-neutral type (Filion et al. 2010; Corrales et al. 2017).

From an evolutionary point of view, although prokaryotes have specialized proteins associated with their DNA, they do not share homology with eukaryotic CAPs (Luijsterburg et al. 2008). In general, evolution of chromatin and diversification of epigenetic mechanisms are suggested to be tightly linked with eukaryotic evolution, from its origin to the transition to multicellularity (Newman 2005; Aravind et al. 2014; Gombar et al. 2014; Penny et al. 2014; Miyamoto et al. 2015; Sebé-Pedrós et al. 2017). Indeed, the Last Eukaryotic Common Ancestor (LECA) is considered to possess the key components of eukaryotic epigenetics, including most histone modification enzymes and some histone mark readers (Aravind et al. 2014). In addition, a current hypothesis on the transition to multicellularity is that complexification of the regulatory genome, via the emergence of repressive chromatin contexts and distal regulatory elements, permitted to generate the cell-type-specific transcriptional programs required for multicellularity (Larroux et al. 2006; Mendoza et al. 2013; Sebé-Pedrós et al. 2016, 2017; Arenas-Mena 2017; Hinman & Cary 2017). Recently, a system-level view of the evolution of chromatin modification machinery was provided by (On et al. 2010). They demonstrated the high conservation of a core of chromatin proteins across four model organisms (human, yeast, fruit fly, and worm), accompanied with diverse lineage-specific innovations. Similarly, a study on the evolution of the DNA damage response network in 47 species found a conserved core of metabolic components (Arcas et al. 2014). Regulatory partners were also present at an early evolutionary age and these steadily diversified over evolution (Arcas et al. 2014).

Here, we investigate the evolutionary relationships of the CAPs studied by (FiliOn et al. 2010; van Bemmel et al. 2013), using homology prediction, gene age inference methods, functional annotations, and protein domain annotations. Taken together, the work provides insight in the conservation of a chromatin landscape across eukaryotes. Our phylogenomic analysis leads us to propose that the chromatin types YELLOW and RED have deep evolutionary roots with many lineage-specific properties. With respect to GREEN chromatin, we provide evidence that some of its associated proteins are undergoing an evolutionary Red Queen process called centromere drive (Malik & Henikoff 2002). Finally, our results support the association between the emergence of BLUE chromatin with its Polycomb proteins, and animal and plant multicellularity.

## Material and Methods

### Data set

Our data set contains all CAPs whose chromatin types have been assigned by (FiliOn et al. 2010; van Bemmel et al. 2013). As a convention throughout the work, a CAP is assigned the color of the chromatin type(s) it binds over more than 10% (fraction of 0.1). The set contains 107 *D. melanogaster* proteins, which include 65 well-characterized CAPs selected to cover a wide range of known chromatin complexes plus 42 previously unknown proteins putatively linked with chromatin. All have also been selected on expressibility in Kc167 cell-lines (derived from *D. melanogaster* embryonic hemocytes). Taking the common assumption that protein function tends to be conserved in homologs across species (Koonin & Galperin 2003), we searched for homologs of CAPs in 53 species, covering 15 prokaryotes, 15 non-metazoan eukaryotes, and 23 metazoa (Supplementary Table 1, Supplementary Figure 1). The selection of species was guided by the quality of their PhylomeDB entry (Huerta-Cepas et al. 2014).

### Homology prediction

All homolog predictions for the set of 107 *D. melanogaster* proteins were extracted using MetaPhOrs (http://orthology.phylomedb.org/) (Pryszcz et al. 2011), a repository of phylogeny-based orthology and paralogy predictions computed through popular homology prediction services: PhylomeDB, Ensembl, EggNOG, OrthoMCL, COG, Fungal Orthogroups, and TreeFam (Tatusov et al. 1997; Chen et al. 2006; Wapinski et al. 2007; Flicek et al. 2008; Ruan et al. 2008; Muller et al. 2010; Huerta-Cepas et al. 2014).

In a first round, we extracted all *D. melanogaster* homology predictions for the 107 CAPs in the other species of interest. We retained only homology hits (i.e orthology and/or paralogy) that had sufficient sequence similarity with the corresponding *D. melanogaster* protein. In all cases, a sequence similarity criterion of 25% and a maximum gap proportion of 60% (i.e. minimum 40% overlap) were applied after Needleman-Wunsch global pairwise alignment with the *D. melanogaster* protein. The maximum gap proportion avoids hits that share very conserved domains in otherwise unconserved sequences. The similarity threshold for homology was chosen to be consistent with knowledge for well-studied proteins, including Polycomb, HP1, SU(VAR)3-9, Sir2, RNA pol, TBP, CTCF, PCNA, SU(HW), BEAF-32 (Klenk et al. 1992; Lanzendörfer et al. 1993; Marsh et al. 1994; Rowlands et al. 1994; Krauss et al. 2006; Lomberk et al. 2006; Whitcomb et al. 2007; Greiss & Gartner 2009; Chia et al. 2010; Schoborg & Labrador 2010; Heger et al. 2013).

The homology prediction of MetaPhOrs is based on searching over half a million pre-computed gene trees. These trees usually focus on subsets of species, for instance, a tree can be restricted to vertebrates only. This may generate false negatives in our first round of homology search, since some species are less likely to appear in trees with *D. melanogaster*. Therefore, a second round of homology search was conducted to cover also the less-studied species as follows. For each protein of a particular organism lacking a hit in the first round, the predicted homologs of the two closest species to that particular organism were used to seed a second search for an ortholog in this organism. For instance, during the first round a homolog of the *D. melanogaster* protein HP6 (HP6_Dme) was found in *D. simulans* as HP6_Dsi, but not in the ant *A. cephalotes*. In the second round, the homology search in *A. cephalotes* was seeded with HP6_Dsi. Then finding an ortholog in *A. cephalotes* points to a candidate homolog of *D. melanogaster* HP6_Dme. We encountered 190 cases of a successful second round of homology search.

Despite the two rounds of homology search, strictly speaking we cannot prove the absence of homologs observed in certain species, as we cannot rule out that it is related to biological and/or technical challenges, such as rapid sequence divergence, limited sequencing depth and/or genome coverage, or the sensitivity of the homology search.

Different amino acid substitution matrices were used to account for different evolutionary distances: Blossum45 to compare with prokaryotes, Blossum62 with eukaryotes, and Blossum80 with metazoa. Finally, we note that instead of *D. melanogaster* Su(var)3-9, the well-characterized human homolog SUV39H2 was used as a seed for homolog search, since this gene and the eukaryotic translation initiation factors eiF2 are fused in *D. melanogaster* (Krauss et al. 2006) and attract false positive hits.

### Gene age inference

The binary vectors of homolog absence/presence of the 107 CAPs for each species were clustered using partitioning around medoids (PAM) (Kaufman & Rousseeuw 1990), with simple matching distance (SMD) as dissimilarity measure, and followed by silhouette optimization. The resulting clustering and age groups are robust, as confirmed by re-runs of PAM and by using the Jaccard distance measure.

Similar to (Arcas et al. 2014), we verify our clustering by independently applying the Dollo parsimony method, which associates gene age to the most recent common ancestor. We relate each gene to the age of the most distant hit, defining 5 age groups: Pre-Eukaryotes, Eukaryotes, Opisthokonta, Metazoa, and Arthropods. For instance, since the most distant homolog of Deformed Wings (DWG) is in the spreading earthmoss *P. patens*, we assign it to Eukaryotes. We confirm that the trends remain unaffected (Supplementary Table 2, Supplementary Figure 2).

Finally, to determine if *D. melanogaster* CAPs are enriched at certain ages, we used ProteinHistorian (Capra et al. 2012) (http://lighthouse.ucsf.edu/proteinhistorian/). ProteinHistorian regroups databases of *D. melanogaster* proteomes with protein age assigned by different methods. We calculated enrichment using five different sets of protein family prediction of the Princeton Protein Orthology Database (Heinicke et al. 2007) (DROME_PPODv4 clustered with OrthoMCL, Multiparanoid, Lens, Jaccard and Panther7) and two different methods (Wagner and Dollo parsimony) to account for the expected differences according to the different phylogenies and data sets (Supplementary Table 3).

### Gene Ontology analysis

We used WebGestalt (Wang et al. 2013) (http://www.webgestalt.org) to search the 107 *D. melanogaster* CAPs for enriched Gene Ontology (GO) terms. 82 of 107 proteins were annotated with GO terms and used for the analysis. We focused on the category Biological Process. Default WebGestalt settings were used to calculate enrichment, p-values were corrected by the False Discovery Rate (FDR) method of Benjamini-Hochberg (BH) and a significance threshold of FDR-corrected p-value < 0.05 was applied. Results were then submitted to REVIGO (Supek et al. 2011) to map the GO terms onto a semantic plane. Using k-means clustering of the semantic (x, y) coordinates, the GO terms were clustered into groups for ease of interpretation. We named these groups manually. To identify trends of GO terms across evolutionary age, we built a background distribution by maintaining the relation (gene, age group) and repeatedly (n=1000) re-assigning to each gene in a random fashion the GO term clusters that were obtained from k-means clustering.

### Reader/Writer/Eraser of histone marks analysis

From the literature, known *D. melanogaster* histone modifiers and histone marks readers were extracted in addition to the ones present in the initial set (Bannister et al. 2001; Cao et al. 2002; Schotta et al. 2002; Byrd & Shearn 2003; Smith et al. 2004; Stabell et al. 2006; Steward et al. 2006; Wysocka et al. 2006; Eissenberg et al. 2007; Larschan et al. 2007; Rudolph et al. 2007; Seum et al. 2007; Srinivasan et al. 2008; Smith et al. 2008; Moore et al. 2010; Rechtsteiner et al. 2010; Wagner & Carpenter 2012). Homologs of these proteins among our species set were searched applying the same method as described in the above section ‘Homology Prediction’.

### Intron/exon extraction and analysis

We extracted genome-wide exon data from Ensembl Biomart (https://www.ensembl.org, v93, and Ensembl Metazoa v40, both released July 2018) for 4 species (*D. melanogaster, A. gambiae, H. sapiens, M. musculus*). Introns were computed by subtracting exons from the coding sequence. For each gene, introns were divided into two groups: the first two 5’ introns, named “first”, and any other introns as “rest”. We verified our method against the *Drosophila* pre-computed intron data of FlyBase and detected no major differences in our results. Note that the pre-computed intron data includes introns in the 5’ UTR. The chromatin type of genes (and their introns) in *Drosophila* was determined by the “color” of their TSS as done in (FiliOn et al. 2010). In the 3 other species, genes were assigned the chromatin color of the ortholog in *D. melanogaster* (using Biomart orthology). Note that a CAP has two colours: one is determined by the chromosomal location of the gene and its TSS, and the other is defined by where the protein binds along the genome together with other CAPs. The evolutionary age of genes was inferred using ProteinHistorian as in the above section ‘Gene age inference’.

### Coding sequences extraction for dN/dS calculation and positive selection tests

For all 107 *D. melanogaster* CAPs, MetaPhOrs was used to retrieve orthologs within ten other Drosophila species (*D. yakuba, D. sechellia, D. pseudoobscura, D. willistoni, D. virilis, D. simulans, D. persimilis, D. erecta, D. ananassae*, and *D. mojavensis*). Using Flybase (http://flybase.org/, version *FB2017_01*, released February 14, 2017), we extracted all corresponding coding sequences (CDS). To avoid different isoforms and different within-species paralogs, only the protein with the highest alignment score to its corresponding *D. melanogaster* protein was retained for each species. Next, with these *Drosophila* species we inferred phylogenetic tree topologies, we estimated dN/dS, and we performed positive selection tests. We elaborate on each of these steps below.

### Sequence alignment and tree topology inference for dN/dS calculation and positive selection tests

To prepare the homology sets for dN/dS calculation and positive selection tests with PAML (Yang 2007), CDSs of each set were multiple-aligned and a tree topology inferred. First, CDSs were translated and multiple aligned with Clustal Omega 2.1 (Chenna et al. 2003) Translation, alignment, cleaning and translation reversion is done with TranslatorX local version (Abascal et al. 2010) (available at http://translatorx.co.uk/), with the following parameters for Gblocks cleaning: ‘−b1=6 −b2=6 −b3=12 −b4=6 −b5=H’ (Castresana 2000). In short, the Gblocks parameters b1 to b4 tune which amino acid (sub)sequences are considered conserved and/or non-conserved. They were chosen to relax cleaning on variable regions and retain diversity. The parameter −b5=H permits to clean sites with gaps in more than half of the sequences, following the recommendation from the PAML documentation to remove such sites. We refer to Gblocks documentation for details.

To account for possible differences between gene trees and species tree, positive selection tests were run on maximum likelihood trees computed from CDS alignments with phyml (GuindOn et al. 2010) and also on *Drosophila* species trees extracted from TimeTree (Kumar et al. 2017) (http://www.timetree.org/). Phyml was run with default parameters to return the topology maximizing the likelihood function.

### dN/dS estimation

From multiple CDS alignments and inferred tree topology (see previous section), PAML fits codon substitution models and estimates both branch length and dN/dS by maximum likelihood. For each of these alignments, a single dN/dS was estimated using Model 0 of codeml included in PAML (Yang 2007). We verified that dN/dS values are similar with the two tree topology inference methods (Supplementary Figure 7).

### Positive selection tests

In order to detect positive selection among amino acid sites and along branches of the *Drosophila* tree, tests were carried out on gene and species trees with codeml from PAML using branch-site codon substitution models (Yang 2007). Since PAML fits models by maximum likelihood, it allows to put constraints on the dN/dS parameter and compare models via their likelihood. Following the approach of “Test 2” (see PAML documentation), we predicted positive selection by comparing Model A to the Null Model. In these models, different constraints can be put on a candidate branch, the so-called foreground branch, and all other branches in the tree, i.e. background branches. Model A allows dN/dS to vary among sites and lineages on the specified foreground branch, thus allowing for positive selection. The Null Model fixes dN/dS to 1 on both foreground and background branches, thus allowing only for neutral selection. This process was automated for all branches in the trees. Finally, for every (Model A, Model Null) pair, likelihood ratio tests (LRT) with Bonferroni correction for multiple testing were applied. The Null model was rejected where the adjusted p-value was < 0.01. Finally, Bayes Empirical Bayes (BEB) calculates the posterior probabilities for sites to be under positive selection when the LRT is significant.

### Protein domain annotation

To search for over-represented domains among GREEN proteins in each of the inferred age clusters, domain annotations were extracted from InterPro database v63 (Finn et al. 2017). DNA-binding domains and their location in D1 proteins from 10 *Drosophila* species were inferred from protein sequence by searching Pfam or Prosite domains using InterProScan v5 (Jones et al. 2014).

## Results

### The *Drosophila* chromatin landscape is dominated by eukaryotic and metazoan-age proteins

Taking the dataset from (Fili On et al. 2010; van Bemmel et al. 2013), we searched for homologs across 53 species and clustered the resulting phylogenetic profile in order to gain insight into the conservation and evolution of the *D. melanogaster* chromatin landscape. The clustering reveals six clusters (Figure 1, left side, I – VI). We associated these clusters to five major age groups: pre-eukaryotic CAPs (I & II), eukaryotic CAPs (III), multicellular plant and metazoan CAPs (IV), metazoan-specific CAPs (V) and arthropod CAPs (VI), with ages assigned on the basis of stable blocks of conserved CAPs in multiple species. For phylogenetic positioning and dates of our five age groups, see Supplementary Figure 1.

We made several major observations on the inferred clusters. We find two dominant clusters, one referring to eukaryotes in general (III) and one specific to metazoans (V), and a third large cluster regarding arthropods (VI), indicating lineage specific diversification. Next, we observe a regular lack of CAPs across evolution, in particular in fungal and parasitic species (for instance *S. pombe* and *S. japonicus*, respectively *Spo* and *Sja* in Figure 1). For fungal species the lack of CAPs may be due to lineage specific divergence, such that we do not detect any homologs, though we cannot rule out lineage specific loss or non-orthologous gene displacement (Koonin et al. 1996). With respect to parasitic species, loss of CAPs is more likely.

**Figure 1.**
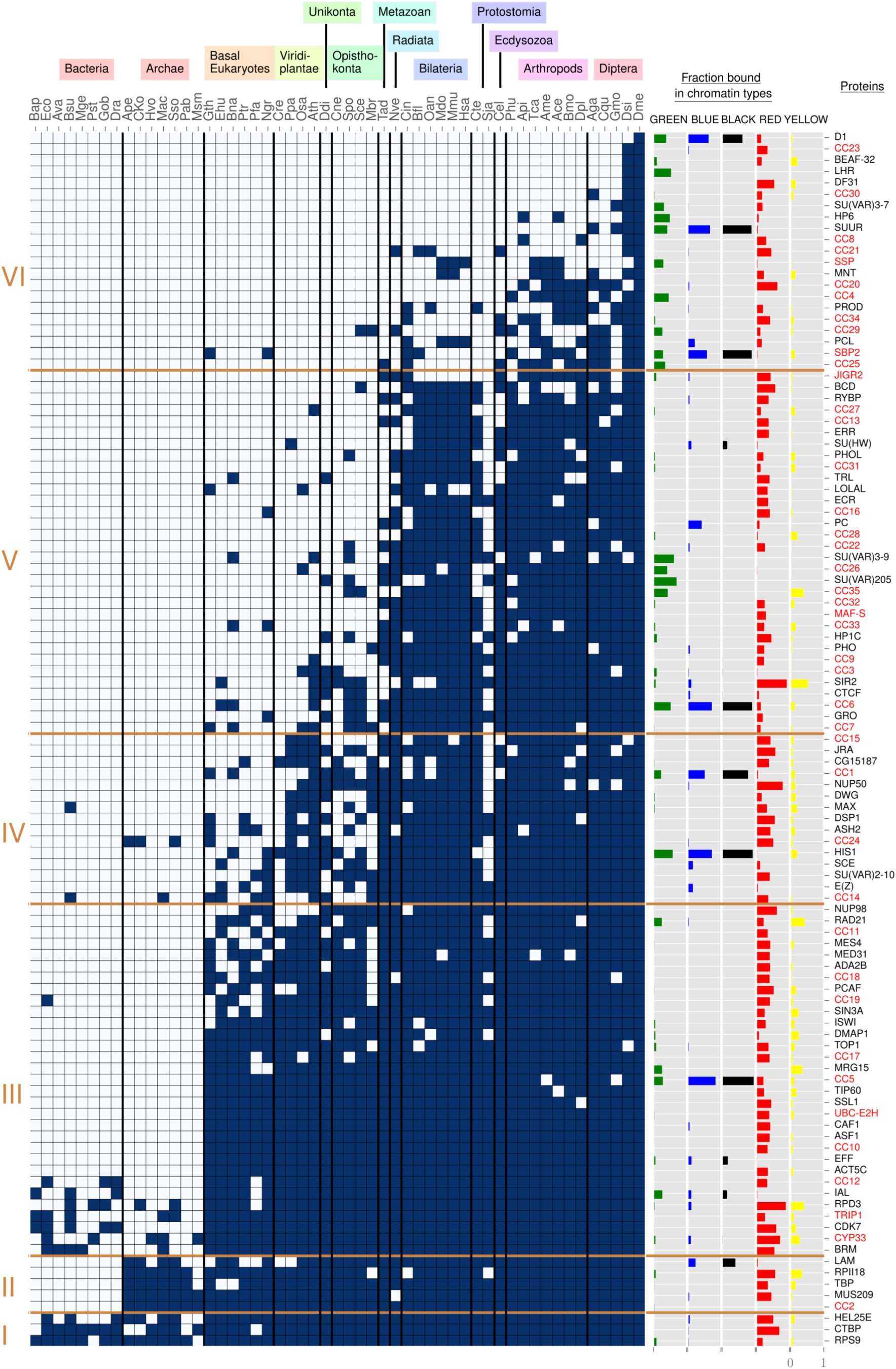
Phylogeneticprofileofchromatin-asociatedproteins. To the left, six protein age clusters resulting from clustering with partitioning around medoids (see Methods for details). They are indicated with Roman numerals (I–VI). On top, 13 species groups are manually defined to aid the reader, three letter codes refer to species names as given in Supplementary Table 1. In the matrix, dark blue rectangles represent the presence of a homolog, grey rectangles its absence. Within each age cluster, rows are ordered from top to bottom by decreasing number of dark blue rectangles. Columns are ordered at the level of species groups by decreasing phylogenetic distance to *D. melanogaster*, with *Drosophila* (Dme) in the rightmost column (see Supplementary Figure 1 for details). Within a species group, columns are arbitrarily ordered. The five columns “Fraction bound in chromatin types” display the fraction of chromatin type (GREEN, BLUE, BLACK, RED, YELLOW) bound by each CAP. To the right, the column “Proteins” contains protein names, with unknown proteins in a red font.

In order to understand what biological functions are present in the data set, we extracted enriched GO terms for the CAPs in the domain ‘Biological Process’ using Webgestalt (Wang et al. 2013). The resulting 37 GO terms were projected onto a semantic plane and clustered to facilitate their interpretation (Figure 2A, see Supplementary Table 4 and Methods for details). As expected for a set of chromatin-associated proteins, we observed highly significant enrichments for the clusters ‘Chromatin organization’ (FDR-corrected p-values ≤1.374e-7) and ‘Regulation of transcription’ (FDR-corrected p-values ≤1.433e-6). The other seven clusters mostly cover basic nuclear processes (‘Transcription’, ‘Protein modification’) and developmental and cell-cycle control (‘Development’, ‘Cell cycle’, and ‘Regulation of cell cycle’). Some GO terms remained difficult to interpret (e.g. number 0 in Figure 2A, ‘Immune system process’), which might reflect the biological process for which a gene or protein was originally annotated.

**Figure 2.**
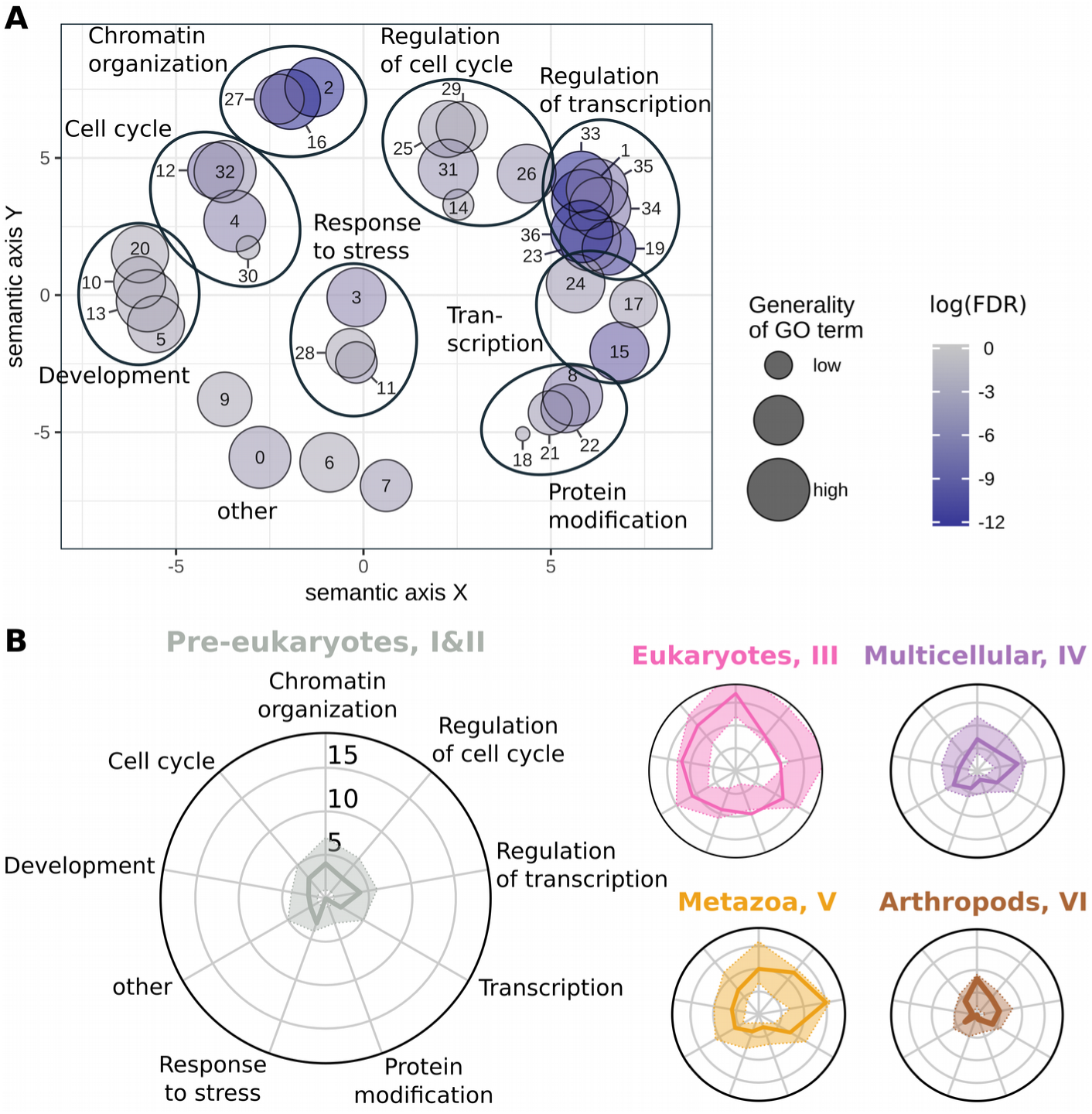
Diversity in functional annotations at evolutionary age groups. **(A)** Enriched GO terms of CAPs projected on a two-dimensional semantic space. The GO terms are clustered into 8 higher-level GO groups as indicated by the large ovals (see Methods for details). The ‘other’ group is left without oval. See Supplementary Table 4 for details. **(B)** For each age group, a radar plot shows the distribution of GO terms from annotated proteins. Axes correspond to the number of CAPs in the higher-level GO groups of panel A. For example, in the leftmost radar plot we observe 4 CAPs in the GO cluster ‘Chromatin organization’. The coloured area around the solid line indicates the 95% boundaries of a randomized background distribution. *Pre-Euk* means pre-eukaryotic gene age (cluster I and II), *Euk* is eukaryotic age (cluster III), *Multicellular* is multicellular plant and metazoan age (cluster IV), *Metazoa* is metazoan age (cluster V), and *Arthropods* is arthropod age (cluster VI).

We then studied the trends in GO terms along the five age groups. With the nine GO term clusters as axes of radar plots, we mapped CAPs through their GO annotations to each axis and visualized the number of CAPs per axis (Figure 2B). Against a randomized background distribution (see Methods for details), we find that most axes do not show over or under-representation of CAPs. Nevertheless, we report several interesting observations. First, ‘Protein modification’ is over-represented amongst eukaryotic CAPs (III) and rather depleted in other age groups. Indeed, 10 of 23 annotated eukaryotic CAPs are involved in histone modifications, especially acetylation (e.g. Rpd3, Ada2b, and Mrg15). Second, eukaryotic CAPs appear depleted of regulatory processes, including cell cycle and transcription. In contrast, metazoan-aged CAPs (V) are enriched for such regulatory processes. This may be interpreted that evolution towards more complex eukaryotic organisms was accompanied by the acquisition of new regulatory interactions. Such a view is consistent with the paradigm that the evolution of increasingly complex transcriptional regulation is one of the key features in (animal) multicellularity, enabling the establishment of precise spatio-temporal patterns of gene expression and regulation (Larroux et al. 2006; Mendoza et al. 2013; Sebé-Pedrós et al. 2016, 2017; Arenas-Mena 2017; Hinman & Cary 2017). Third, the youngest age group of arthropod CAPs is enriched in chromatin organization. We will provide an explanation for this observation on the basis of fast evolving GREEN proteins in later sections. Finally, we strengthened the importance of the eukaryote cluster (III) by independent age enrichment tests against the full *D. melanogaster* proteome, with age assigned to each protein by means of Dollo and Wagner parsimony (Csurös 2010). Indeed, we find that CAPs are significantly enriched in genes that date back to the origin of eukaryotes (Fisher’s exact test, see Supplementary Table 3 for details).

In summary, many CAPs appear to have been established early in eukaryotic evolution, as was also reported in (Aravind et al. 2014). And the evolution of regulatory proteins may have been especially important during the evolution toward more complex multicellular organisms. In the next sections, we assess the conservation of the *D. melanogaster* chromatin landscape in eukaryotes and we highlight three major dynamics in chromatin evolution.

### YELLOW and RED chromatin both date back to early eukaryotic evolution

Of the five chromatin types, YELLOW and RED are the two euchromatic types, associated to transcriptionally active regions in the genome. One of the key differences between genes in YELLOW and RED is their expression pattern across embryonic stages and tissues (Fili On et al. 2010). YELLOW genes are broadly expressed and have predominantly housekeeping functions, while RED ones have specific expression patterns and are strongly related to developmental processes.

We formulated two contrasting explanations for the evolutionary history of euchromatin and its associated proteins, and assessed the evidence for both. First, YELLOW and RED may derive from a single ancestral euchromatic type. RED evolved from this common type to address the challenges of multicellular life and development, and as such should consist of relatively young CAPs. On the other hand, developmental processes are built upon the cell’s machinery to respond in a timely and proportionate fashion to environmental cues. This is an inherent (old) feature of any cell and one could speculate it is qualitatively different from regulating housekeeping genes. Thus our second hypothesis is that YELLOW and RED address a functional difference in the regulation of gene expression, which dates (at least) from the origin of the eukaryotic cell.

Below we present three lines of inquiry. Taken together, these suggest that both YELLOW and RED chromatin date (at least) back to LECA. This interpretation disfavours the explanation that RED specifically evolved for regulating development.

#### Young YELLOW and RED CAPs are more specialized than old ones

We asked if the phylogenetic profile of CAPs supports one of the explanations introduced above. Thus we examined CAPs and their coverage in YELLOW and RED over evolutionary time (Figure 1). First, we took for each protein its coverage in YELLOW and RED (Figure 1). For instance, NUP98 has a low coverage in YELLOW (2.5%) and high in RED (63.1%). We found that, on average, proteins in the pre-eukarotic and eukaryotic age clusters bind YELLOW and RED more abundantly than younger proteins (Figure 3A). For instance, the 32 metazoan-age CAPs bind less in RED genomic regions than the 8 pre-eukaryotic ones (median coverage 0.24 against 0.41, p-value = 0.105, one-sided Mann-Whitney-U test). This most likely reflects “old” general transcriptional machinery and “young” regulators of gene expression. Second, we classified each protein as present in a given chromatin colour if its coverage > 0.1. In this manner, the protein mentioned above, NUP98, is considered only bound in RED. With this categorization, old proteins more often associate with both YELLOW and RED, while younger ones appear to be more specialized to one of the two types (Figure 3B). If there used to be a single ancestral euchromatic type, one could indeed expect that older proteins discriminate less between YELLOW and RED. Yet, it may also reflect a degree of shared nuclear machinery (e.g. RNA polymerase RPII18), which is consistent with the second hypothesis.

**Figure 3.**
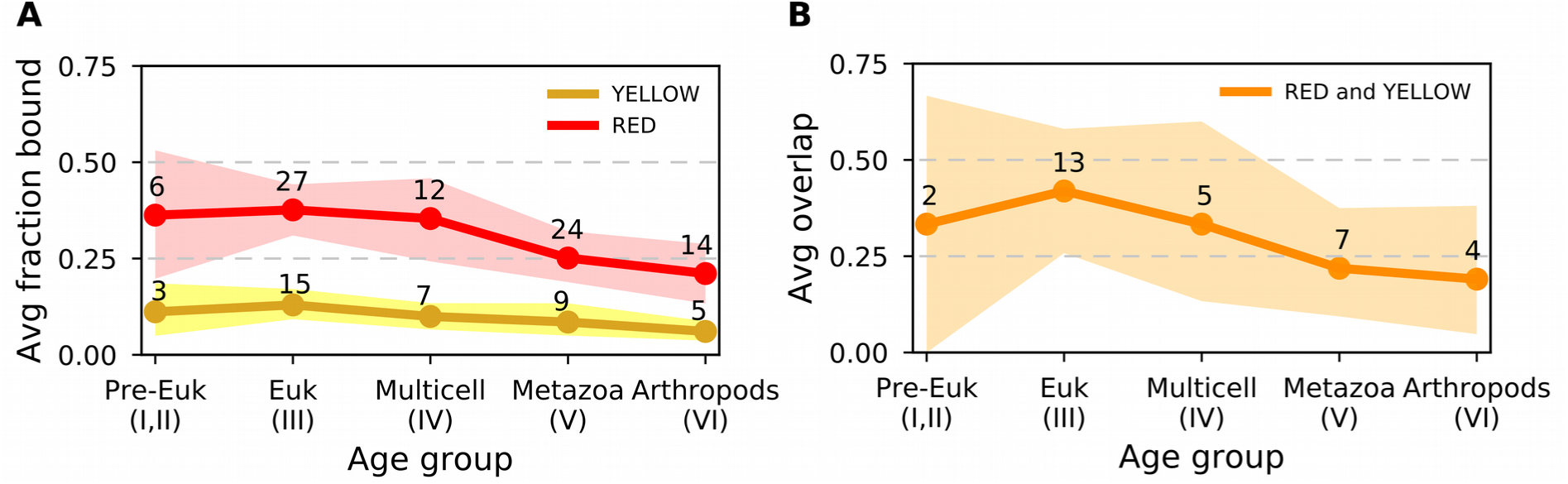
Average fraction of genome bound by proteins over evolutionary age groups. **(A)** Per age group, the average fraction of YELLOW and RED chromatin to which CAPs bind, with 95% confidence intervals obtained by bootstrap analysis. The average is calculated on the ‘raw’ fraction bound as displayed in Fig. 1. In addition, points annotated with numbers, indicate the number of proteins per age group classified as bound to a given chromatin type (fraction for a chromatin type > 0.1). **(B)** Average proportion of CAPs of each evolutionary age group that bind both YELLOW and RED (fraction for both chromatin types > 0.1). Again, 95% confidence intervals were obtained by bootstrap analysis, and annotated points are number of proteins bound per age group. In both panels, the bootstrap procedure was as follows: for each age group, say of size *n*, we resample *n* proteins for 1000 times. For panel A, we compute each time the average fraction of YELLOW and RED bound by the resampled proteins, while for panel B, we count the CAPs that bind both chromatin types > 0.1. See Fig. 1 “Fraction bound in chromatin types” for the fraction of chromatin bound by individual proteins.

#### Gene structure in YELLOW and RED is lineage specific

Next, we explored the intron/exon structure of genes in YELLOW and RED genomic regions. In *Drosophila*, housekeeping genes (in YELLOW) have few, relatively short introns, and developmental genes (mostly in RED) start with long 5’ introns and have short ones at the 3’ end (Corrales et al. 2017). We wondered if genes with long 5’ introns mostly date to the origin of multicellularity, and if this feature is also found among other multicellular organisms.

A general feature of gene architecture in eukaryotes is that first introns tend to be longer than the ones that follow (Bradnam & Korf 2008). Indeed, long introns close to the start of a gene are thought to harbour regulatory elements (Chung et al. 2006; Bradnam & Korf 2008; Cenik et al. 2010). We confirmed that genes in *Drosophila* and their orthologs in the mosquito *A. gambiae*, human, and mouse have longer first introns (Supplementary Figure 3), and this remains valid if we subdivide the genes by chromatin colour (Figure 4A) and by estimated evolutionary age (FDR-corrected Mann-Whitney-U tests, Supplementary Table 5). Moreover, in all four species, RED genes tend to have longer first introns than YELLOW genes (Figure 4B). However, insects and mammals differ when we subdivide YELLOW and RED genes by evolutionary age (Supplementary Table 5). Fly and mosquito suggest that differences in the first introns trace back to the origin of LECA (but not before). Human and mouse, on the other hand, display only few significant differences between YELLOW and RED first introns over evolutionary age. Overall, we do not find a clear signal that RED’s typical gene structure of long 5’ introns is a (general) evolutionary response to regulate developmental processes. Instead we propose that gene structure in YELLOW and RED is dominated by lineage-specific features.

**Figure 4.**
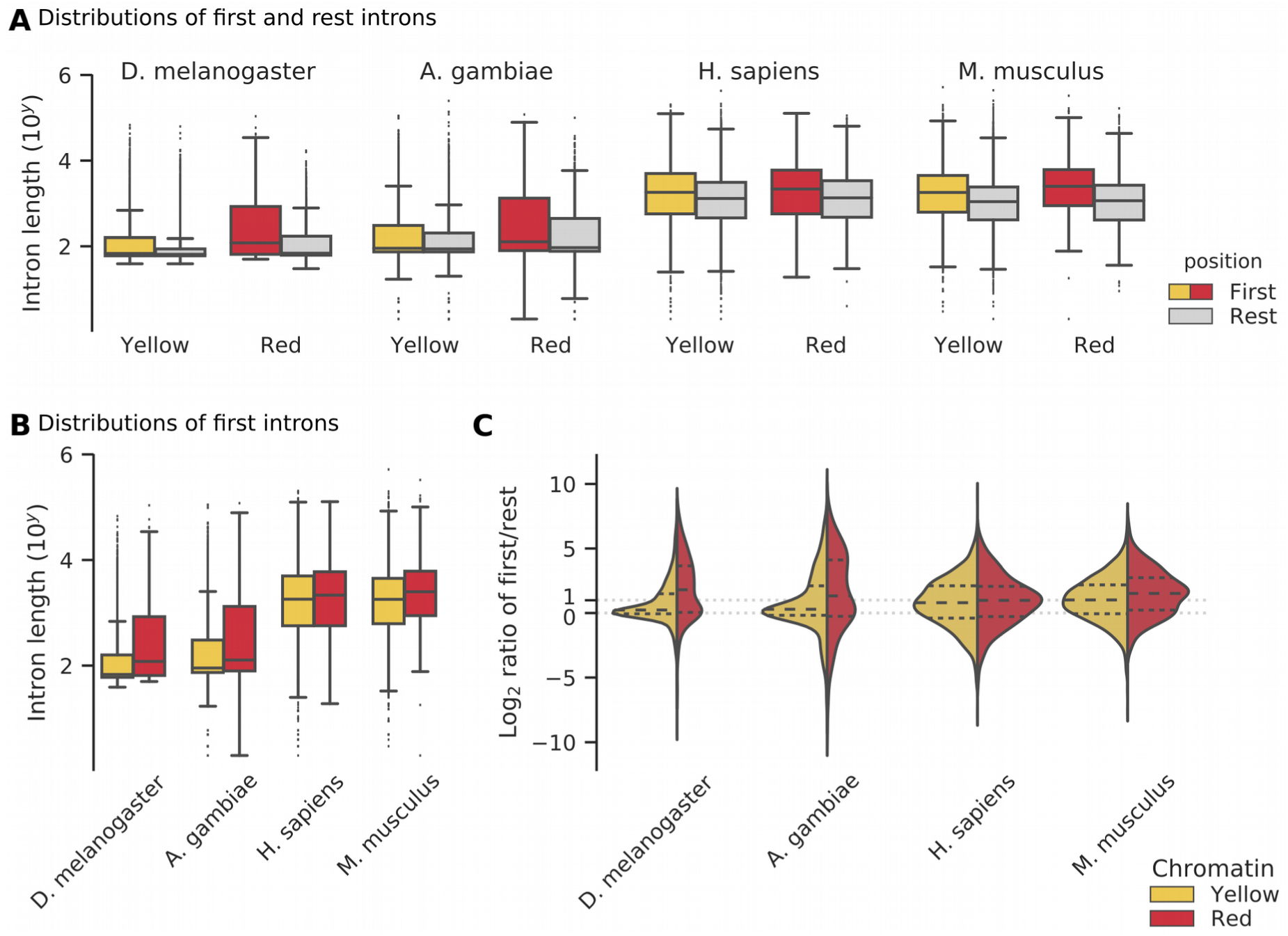
Intron structure of genes in YELLOW and RED genomic regions. **(A)** Length distributions of the first two 5’ introns and the rest introns in two insects and two mammals. The distributions are subdivided by chromatin type (Yellow, Red). Note the logarithmic scale of the y-axis and the large number of long outliers for fruit fly and mosquito. **(B)** Direct comparison of the distributions of first intron lengths in YELLOW and RED genomic regions. **(C)** Intron length ratio distributions. Plots are split into a YELLOW (left) and RED (right) distribution for each species (see panel A). Each half shows log2 ratios of the following: per gene we divide the median length of the first two 5’ introns by the median length of other introns (toward the 3’ end). Each violin plot has quartiles marked inside the coloured area at fractions of 0.25, 0.50, and 0.75. We indicate the equal ratio of intron lengths (ratio = 0.0) and the ratio of 2-times as long first introns (ratio = 1.0).

#### Gene structure in *Drosophila* YELLOW and RED is more variable than a bulk comparison suggested

Until now we compared first introns and other introns ignoring that they are part of the same genes. We calculated the ratio of first against others on a per gene basis, and we repeated our analysis. Surprisingly, we find that the four species have a substantial subset of genes that have smaller first (5’) introns compared to their 3’ introns (Figure 4C, log2-ratio < 0.0). For YELLOW and RED genes in fly and mosquito, the intron ratio distribution is composed of two kinds of genes (Figure 4C). The central peak at log2-ratio ∼0.0 indicates a subset of genes with equal intron sizes along the coding sequence, and second subset of genes with long first introns is signalled by a ‘shoulder’ at log2-ratio > 1.0. We do not observe bimodal distributions for human and mouse (Figure 4C).

We wondered what kind of genes compose the sub-distributions in *Drosophila.* We find that YELLOW genes with long first introns (log2-ratio >1.0) are relatively enriched for GO terms relating to development in comparison to YELLOW genes with equal-sized introns (log2-ratio ∼0.0) (WebGestalt GO analysis, FDR < 0.004, see Supplementary Table 6). Vice versa, YELLOW genes with equal-sized introns are relatively enriched for RNA and DNA-related processes. For RED genes of either subset, we do not detect any significant GO term enrichment. We conclude that while the trend is for YELLOW to contain housekeeping genes, it also has development-related genes. We come back to this in the Discussion.

#### Histone modifications in YELLOW and RED are ancient and shared by both euchromatin types

Our third analysis addresses histone modifications associated to euchromatin. Filion et al. focused on H3K4me3, which is found in close vicinity to active Transcription Start Sites (TSS), and H3K36me3, which marks transcribed exons. Though reported slightly differently in their original paper, we know now that both YELLOW and RED are marked by both tri-methylations (Corrales et al. 2017). We summarize here, what is known about the evolutionary conservation of the two marks.

Given that the last eukaryotic common ancestor (LECA) had a lysine (K) at the amino acid positions indicated by H3K4 and H3K36 (Aravind et al. 2014), we created a phylogenetic profile of histone-related proteins across 53 species, similar to the profile that we made for CAPs (Supplementary Figure 4). We focused on three classes of proteins: *writers* that modify the histone (i.e. methylation), *readers* that interpret the mark, and *erasers* that remove the mark. We identified the first putative writer for both H3 lysine marks in one basal eukaryote (*P. tricornutum*) and three viridiplantae (*P. patens, O. sativa* and *A. thaliana*). And we found a H3K4me3 reader and H3K36me3 eraser in four basal eukaryotes (*G. theta,E.huxleyi, B. natans*, and *P. tricornutum*). Next, we examined literature evidence for the histone modifications themselves. Studies in yeasts, plants, and *Capsaspora owczarzaki* (a close unicellular relative of metazoa) reveal abundant use of both H3K4me3 and H3K36me3 (Bernstein et al. 2002; Suzuki et al. 2016; Roudier et al. 2011; Sebé-Pedrós et al. 2016). In addition, basal unicellular eukaryotes such as *Tetrahymena, Euglena, Stylonychia*, and *Trichomonas* use H3K4me3, though evidence for H3K36me3 is currently lacking (Garcia et al. 2007; Postberg et al. 2010). Overall, these observations suggest H3K4 and H3K36 methylation are ancient, functional epigenetic marks. In the Discussion, we place these results in the context of euchromatin evolution.

### GREEN evolves fast and expanded in a lineage-specific way in *Drosophila*

GREEN chromatin is best characterized as constitutive, classic heterochromatin, and encompasses regions with high content in repetitive DNA and transposable elements (Sun et al. 1997; Filion et al. 2010). It is marked by HP1, a protein family that is involved in chromatin packaging and that binds di-and trimethylated histone H3 (H3K9me2/3) (Bannister et al. 2001). Classic proteins linked with HP1 heterochromatin are conserved (Saksouk et al. 2015) and indeed we find HP1 (SU(VAR)205 in Fig. 1), HP1c, and SU(VAR)3-9 across metazoa (cluster IV). Yet, 11 GREEN proteins, from a total of 25 in the whole dataset, are assigned to the arthropod cluster, the youngest gene cluster VI. Thus, as opposed to YELLOW and RED, the fraction of proteins bound in GREEN increases through evolutionary times (Figure 5B). At first view, this observation is paradoxical, since GREEN proteins are involved in genome integrity, in particular centromere maintenance. One expects to find them conserved across metazoa.

**Figure 5.**
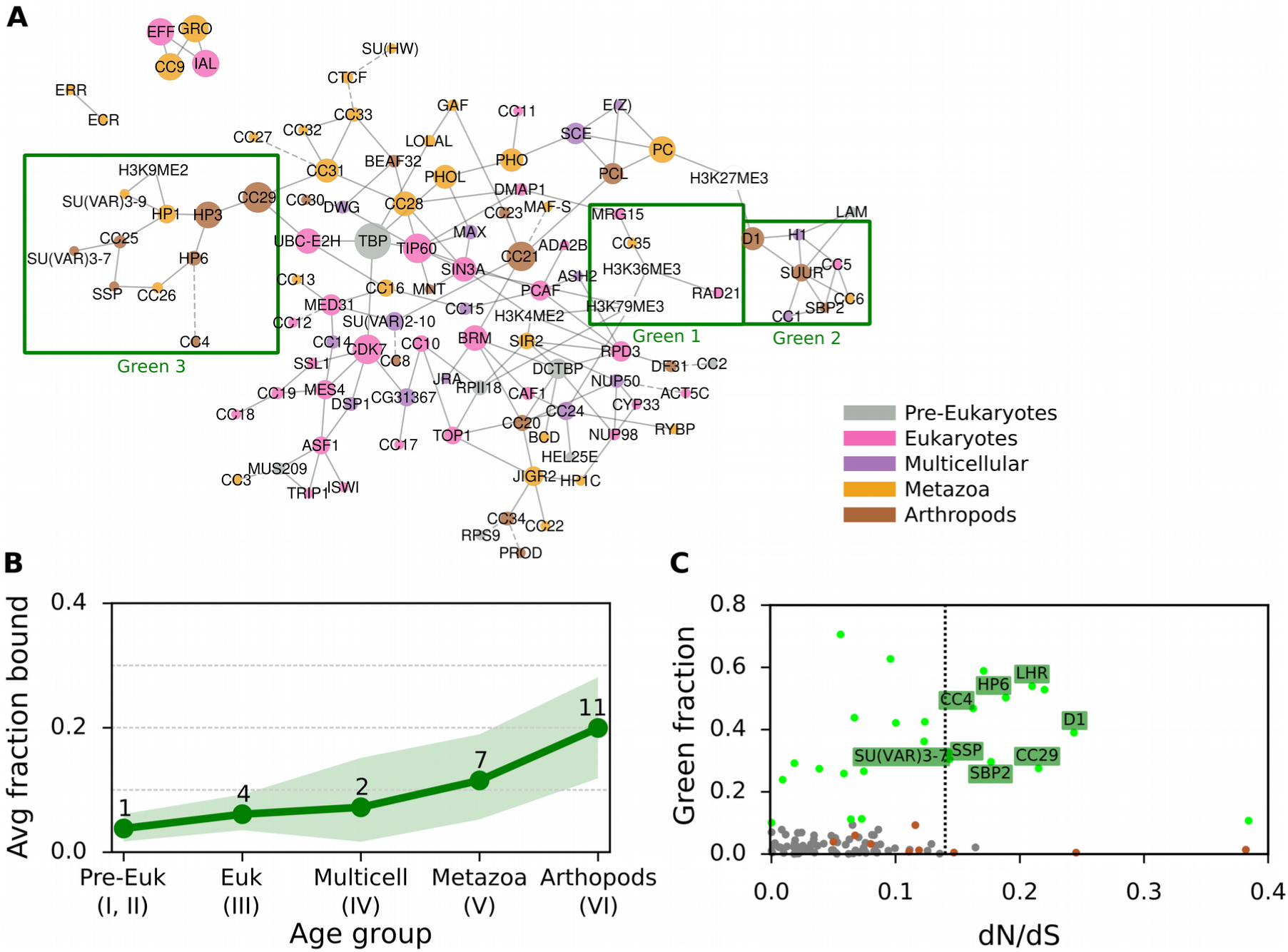
GREEN-associated proteins over evolutionary age and their dN/dS ratio. **(A)** Bayesian network of CAPs adapted from (van Bemmel et al. 2013), with each CAP coloured by age. Proteins and histone marks are connected by a solid line if the predicted interaction has a confidence score of at least 70%. Dashed lines indicate the highest scoring interaction for proteins with all confidence scores below 70%. Three GREEN chromatin subnetworks are highlighted with a green box. **(B)** Average fraction of GREEN chromatin bound by CAPs of each evolutionary age group, with 95% confidence intervals obtained through bootstrap analysis. See Figure 1 “Fraction bound in chromatin types” for the individual proteins and their fraction of chromatin bound and Figure 3 for details on the bootstrap method. **(C)** The ratio of non-synonymous over synonymous aminoacid mutations (dN/dS) against the GREEN fraction bound for each of the 107 CAPs. The dotted vertical line divides CAPs with dN/dS > 0.135 (15% higher of distribution) from the rest. Chromatin proteins are grey dots, with GREEN ones as green dots. GREEN Arthropod Cluster proteins (GAPs) are labelled with their names, while other proteins of the Arthropod Cluster are brown dots.

In a first step, we explored the conservation of GREEN chromatin proteins in the context of a previously established *Drosophila* chromatin protein network. In a pioneering effort, (van Bemmel et al. 2013) applied Bayesian network inference on binding profiles of CAPs to model interactions among chromatin components. In this model, GREEN is the only chromatin type to be divided into multiple regions of the network (Figure 5A), which has led to the suggestion that GREEN chromatin is decomposable into three distinct subtypes (van Bemmel et al. 2013).

We propose this fragmentation to be linked to gene age. In the network, Region 1 contains 3 proteins, RAD21, MRG15, and CC35, that bind both GREEN and YELLOW chromatin. They belong to the oldest group of GREEN proteins. RAD21 and MRG15 are found across eukaryotes (cluster III), while CC35 is predicted to be of metazoan origins (cluster IV). Region 2 consists of proteins of all age clusters, from eukaryotes to arthropods, marking the two heterochromatin types (GREEN, BLUE) and BLACK. The region is organized around SUUR, a key player in chromatin silencing on polytene chromosomes (Makunin et al. 2002). Finally, region 3 contains mostly young GREEN proteins from the arthropod age group, organized around two metazoan proteins, HP1 and SU(VAR)3-9. Matching the three regions to the protein age clusters, we find that regions 2 and 3 are strongly involved in the specific expansion of young GREEN proteins in *Drosophila*. Moreover, their peripheral location in the chromatin network compared to region 1 is consistent with this explanation (Zhang et al. 2015).

### *D1 chromosomal protein* may evolve under the centromere drive model

We asked if poor conservation of many GREEN proteins may be due to the fact that they are fast evolving, which would lead to the rapid divergence of homologs. We estimated dN/dS, the ratio of non-synonymous nucleotide substitutions versus synonymous substitutions among different *Drosophila* species for all CAPs (Figure 5C). Under neutral evolution, non-synonymous substitutions and synonymous substitutions occur with the same probabilities and dN/dS ∼ 1. If positively selected, amino acids change rapidly and dN/dS > 1. On the other hand, under purifying selection amino acid variation is reduced and results in dN/dS < 1. The ratio averaged over all sites and all lineages is however almost never > 1, since positive selection is unlikely to affect all sites over long periods of time. Our analysis revealed that GREEN CAPs from the arthropod cluster (GREEN Arthropod Cluster, GAC) show significantly more elevated dN/dS than other CAPs (8 GACs among a total of 16 CAPs with elevated dN/dS, p-value = 0.0054, Pearson’s Chi-squared test with Yates continuity correction; moderate effect size, Cramer’s V = 0.237). We verified that the over-representation of GACs is not due to young age or protein domain architecture. First, using a logistic regression, we found that dN/dS explains being GREEN (p-value = 0.020) and age does not (p-value = 0.188). Second, we took into consideration that dN/dS is influenced by structural properties of a protein (i.e. unordered protein regions allow for higher dN/dS, while protein domains usually show low dN/dS). We computed dN/dS for 100 randomly selected proteins with similar domain compositions as the young GREEN proteins (Supplementary Figure 5 and 6). With some exceptions, the random selection of proteins have a similar dN/dS as the majority of the CAPs. We concluded that also protein domain composition does not explain the elevated dN/dS.

**Figure 6.**
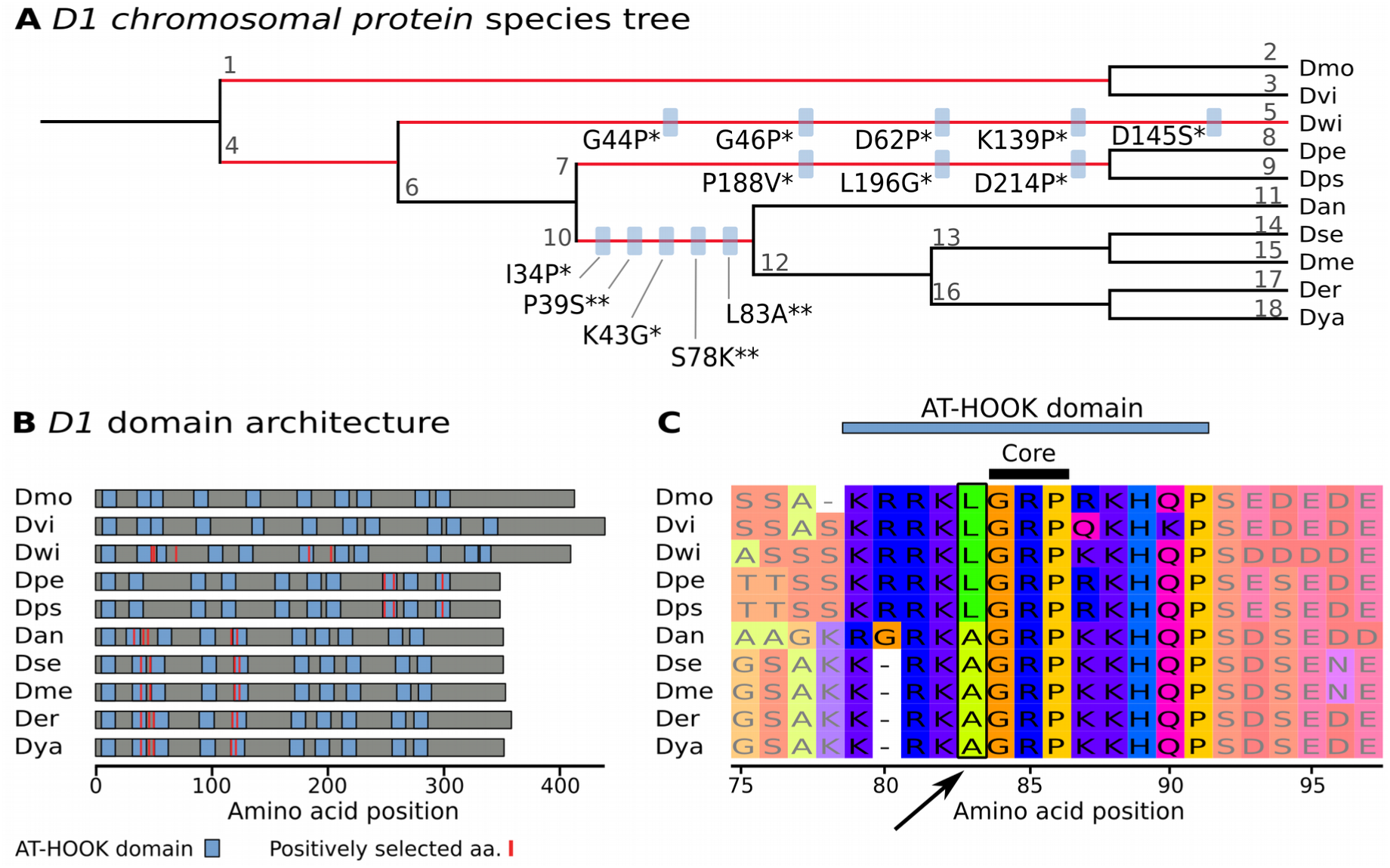
*D1 Chromosomal protein* is undergoing positive selection at AT-HOOK domains. **(A)** Drosophilid species tree (with arbitrary branch lengths). The five branches with positive selection events are highlighted in red (p< 0.01, Bonferroni correction). On branches with more than one positively selected site, blue boxes indicate the amino acid substitution under positive selection, with the significance given as posterior probability of dN/dS > 1 (* for Pr > 0.95, ** for Pr > 0.99). For instance, L83A indicates the substitution of lysine with alanine at position 83. The *Drosophila* species are indicated by 3 letter abbreviations: Dmo is *D. mojavensis*, Dvi is *D. virilis*, Dwi is *D. willistoni*, Dpe is *D. persimilis*, Dps is *D. pseudoobscura*, Dan is *D. ananassae,* Dse is *D. sechellia*, Dme is *D. melanogaster*, Der is *D. erecta*, and Dya is *D. yakuba.* Each branch is identified with a number that links to details on positive selection tests (Supplementary Table 7). **(B)** Domain architecture of D1 proteins. Per species, protein length is given by grey horizontal bars. AT-HOOK domains are represented by blue boxes and positively selected amino acids (aa) are displayed as red vertical lines. **(C)** An example of positive selection in an AT-HOOK domain. The multiple alignment of D1 is zoomed in on the region 75–97aa, showing a change of L83 directly in front of the core motif. Visualisation is done with MSAViewer (http://msa.biojs.net/app/).

Next, we asked if the 8 GAC candidates evolve under relaxed selective constraint or under positive selection. In particular, we wondered if they fit the centromere-drive model proposed in (Henikoff & Malik 2002; Brown & O’Neill 2014). In this model, some heterochromatin proteins evolve under positive selection to suppress the deleterious effect of genetic drive in meiosis. This genetic drive is the consequence of a selfish behavior of chromosomes, which compete for preferential transmission in female meiosis by increasing affinity for microtubule attachment. Chromosomes with more satellite DNA sequences gain an advantage, if heterochromatin proteins involved in recruitment of microtubules do not correct the bias by changing binding specificity. If a centromere drive is left unchecked, it breaks meiotic parity and has a deleterious effect on fitness both at the organism level and at the species level. Chromatin proteins repressing the drive must therefore contain both a role in binding satellite DNA and a role in recruitment of other heterochromatic or centromere proteins.

Of the 8 GACs candidates, LHR (HP3) and HP6 have been proposed to be evolving under this model (Brideau et al. 2006; Ross et al. 2013). We carried out a positive selection test under a branch-site model and found recurrent positive selection for D1. D1 presents the features of heterochromatin proteins evolving through centromere-drive: it is capable of binding satellite DNA and is involved in heterochromatin propagation (Levinger & Varshavsky 1982) To the best of our knowledge, it has not been previously reported as a centromere drive protein. We also speculate that CC29 is a potential candidate. Although we have not been able to detect positive selection using the branch site model, CC29 has DNA binding domains, shows elevated dN/dS, and is part of a centromeric complex with HMR and LHR (Thomae et al. 2013).

For a better characterization of how positive selection is affecting D1 and to corroborate the hypothesis that it is involved in the centromere drive, we investigated more closely at which amino acids positive selection took place. We detected that positively selected sites are within or close to AT-HOOK domains (Figure 6A, B). AT-HOOK domains enable D1 to bind to DNA: the domain is organized around a so-called GRP core, which is able to insert itself into the minor groove of DNA (Aravind & Landsman 1998). Many negatively charged amino acids around this core are then involved in DNA-protein interactions. *Drosophila* species have nine to eleven copies of AT-HOOK in D1 (Figure 6B). Moreover, their locations in sequences vary between species (Figure 6B), highlighting domain-level differences in D1 proteins amongst *Drosophila,* possibly related to DNA binding specificity. As an example of a positively selected amino acid in an AT-HOOK motif, Leucine 83 is replaced by an Alanine directly before the GRP core (Figure 6C). We verified that positively selected sites are equivalent between the two tree topology inference methods, i.e species tree and gene tree (Supplementary Figure 7 and 8, Supplementary Table 7 and 8). In summary, D1 shows strong signs of evolving under positive selection in *Drosophila* and we surmise that it tunes the specificity of its DNA-binding motifs to counterbalance fast-evolving satellite DNA.

### Recent GREEN proteins associate with the expansion of the BESS protein domain in drosophilids

We studied the evolution of the GREEN proteins that lacked signs of positive selection. Notably, in the *Drosophila* genus, the HP1 family has been demonstrated to present little evidence of positive selection. Nevertheless, this protein family is numerous with about 25 members, of which only four are conserved across a large number of drosophilids, and others are evolutionarily restricted to particular *Drosophila* species (Levine et al. 2012). This diversification of the HP1 family is thought to be a lineage-specific expansion driven by karyotype evolution, where events of chromosome rearrangements (fusion/fission) correlate with losses and gains of HP1 proteins (Levine et al. 2012). We explored if other GREEN-associated proteins showed signs of lineage-specific expansions in *Drosophila*.

By studying protein domains, we found evidence that a subset of young GREEN proteins are part of the family of proteins with BESS domains that is expanding in the *Drosophila* lineage. BESS domains direct protein-protein interactions, including with itself. Among all known proteins (not just the ones in our data set) with an inferred BESS domain (InterPro identifier IPR004210), more than 80% are restricted to insects and more than 50% are restricted to diptera. A comparison among drosophilids has shown that the BESS domain family expanded through duplications in a lineage-specific way approximately 40 million years ago (Shukla et al. 2014). In our dataset, five of 107 proteins have a BESS domain (SU(VAR)3-7, LHR, BEAF-32, CC20, and CC25). They are all found in the arthropod cluster (VI), and with the exception of CC20, they are GREEN-associated. Therefore, we propose that these GREEN CAPs evolve rapidly through lineage-specific expansion. And we speculate that BESS domains have a role to play in directing protein-protein interactions in GREEN chromatin in *Drosophila*.

### BLUE is related to the origin of multicellularity

Central in BLUE chromatin are the Polycomb group (PcG) proteins, which are recruited to Polycomb Response Elements (PREs) to silence specific target genes during development, such as Hox genes. PcG proteins form two multiprotein complexes, PRC1 and PRC2. Their catalytic signatures are well-characterized; PRC2 trimethylates histone H3K27 into H3K27me3; this modified histone is bound by PRC1, which in turn ubiquitylates histone H2A. Extensive study on the evolution and conservation of PRC1 and PRC2 has suggested that expansion and diversification of PcG proteins contributed to the complexity of multicellular organisms (Trojer & Reinberg 2006; Whitcomb et al. 2007; Köhler & Villar 2008; Gombar et al. 2014).

In this study, the PcG proteins are represented by the main components of PRC2, namely E(Z) and PCL, and PRC1, with SCE and PC, in addition to three PRE-binders, respectively PHO, LOLAL, and PHOL. PRE-binders are found in RED chromatin, though, as they trigger the transition from active developmentally controlled chromatin to the PcG repressed state. Of the PcG proteins, the oldest ones that lay down key heterochromatin histone marks, are found in the multicellular cluster (IV). They are the writers E(Z) and SCE, which, respectively, tri-methylate H3K27 and ubiquitinate H3K118. Another key BLUE protein, PC, which reads H3K27me3 marks, is metazoan (Cluster V). This is in support of the hypothesis that PRC1, which contains PC, is younger than PRC2. Summarizing, both complexes are conserved across metazoans, suggesting the repression mediated by the PcG proteins as described above, was established at the origins of animal multicellularity (Whitcomb et al. 2007).

Several BLUE proteins are found in cluster II and III, and thus are older than PcG proteins. We mention the three most prominent ones: EFF, IAL, and LAM. All three are conserved in all eukaryotes, with functions unrelated to Polycomb-controlled repression. EFF is involved in protein ubiquitination and degradation, and is suggested to have a general role in chromatin organization (Cipressa & Cenci 2013). IAL is mainly involved in mitosis (Adams et al. 2001) and LAM recruits chromatin to the nuclear envelope (Gruenbaum et al. 1988). As observed by Filion et al., these are general heterochomatic proteins recruited in GREEN, BLUE, and BLACK chromatin to form a repressed state (Figure 1).

## Discussion

We have presented an integrated view of the evolution and conservation of a chromatin-associated proteome across eukaryotes. The creation and analysis of a phylogenetic profile of protein presence/absence across >50 species resulted in three major findings. We find initial evidence in support of the idea that YELLOW and RED chromatin have their roots in early eukaryotic evolution. Second, GREEN-associated proteins were found to be relatively specific to arthopods, or even restricted to dipterans. We connected two evolutionary processes to this observation, namely a Red Queen type of evolution due to centromere drive, and lineage-specific expansion of proteins with BESS domains. Finally, our analysis of BLUE chromatin confirmed existing hypotheses on the importance of Polycomb repressive proteins for the evolutionary success of multicellular life forms. The fifth and last chromatin type, BLACK, has not been addressed in this work. Even if it covers approximately half the genome, it is hard to interpret because it is mechanistically poorly understood and its proteins overlap strongly with those of BLUE chromatin.

### Caveats

To place these results in context, we discuss some critical points of our study. First of all, there is currently no complete list of proteins associated to chromatin. The proteome established by (Filion et al. 2010; van Bemmel et al. 2013) is the most complete one available for metazoans, yet it may have biases that colour our results. The original data set of Filion *et al.* (2010) was mainly based on known chromatin-related processes, such as transcription, regulation of transcription, histone modifiers, nucleosome remodellers, and structural components. Selecting only known proteins necessarily introduces a bias in the data set; these proteins may not uniformly address all chromatin-related dynamics. This was recognized in (van Bemmel et al. 2013). Indeed, they doubled the number of proteins in the data set by scanning the genome for potential chromatin-associating genes and experimentally establishing their chromatin binding pattern. Many of the newly added proteins with a clear association to chromatin were novel, unknown components (e.g. any CC-named protein in the data set is a new Chromatin Component). This was an important effort to create a more comprehensive chromatin proteome. Yet, biases remain due to potential experimental and technical issues. For instance, the genome scan may exclude bona fide CAPs due to the search criteria. Thus one has to keep in mind that the evolutionary trends that we detect can be influenced by a bias in the data set. On a more philosophical note, the proteome used in this study is the current state of knowledge in the field, and as this body of work will be improved upon in the future, so will its interpretation.

Second, the evolutionary view on an epigenetic landscape that we have provided here is, of course, restricted in the sense that it is defined explicitly from a *D. melanogaster* angle. Notably, the *Drosophila* genome is particular, as it appears to lack DNA methylation and is known for an original mechanism of telomere maintenance by specialized non-LTR retrotransposons (Pardue & DeBaryshe 1999). Also, the homologs of *D. melanogaster* CAPs in other species do not necessarily share the same interactions and global assembly to form similar chromatin types. Indeed, in distant species that are separated by more evolutionary time, they are more likely to be functionally different. To counter such false positives, we used a strict similarity cut-off for all protein-protein comparisons. The cut-off indeed helped us to reject functional homology prediction. For instance, it did not accept the *A. thaliana* HP1 homolog, LHP1, which appears to function both in a “classical” HP1-fashion and as a PcG protein (Zhang et al. 2007). Nevertheless, we cannot exclude that even if sequences and domains are very similar, the exact role in chromatin organization may be different.

Finally, as with most proposals of a specific unfolding of the evolutionary process, we note that there is an element of speculation present. Especially regarding our results on the evolution of the euchromatic types YELLOW and RED, we stress that our hypotheses are best seen as initial guides to focus the analysis. We do not claim, of course, that they are the only reasonable explanations regarding the evolution of euchromatin. Indeed, as we explored the ratio of first/other introns, we observed subsets of genes in YELLOW and RED that do not easily fit the two proposed explanations. One alternative is that we observe limitations of the *Drosophila* cell-line used for the experiments. A cell-line is not in the same (micro)environment as a normal *Drosophila* cell and the dynamics of CAPs in the nucleus may be altered. In addition, not every developmental gene may always be marked in RED. While genes that should remain silenced, will be packed into Polycomb-related BLUE heterochromatin, a third category of developmental genes might exist that is marked by YELLOW.

### Histone modifications, gene regulation, and the origins of multicellularity

The evolution of (animal) multicellularity is one of the major transitions in evolution. Within the area of (epi)genomics, it has been hypothesized that complexification of chromatin states and in particular the emergence of distinct heterochromatin states lay at the origin of multicellular life (Sebé-Pedrós et al. 2016; Hinman & Cary 2017). For instance, general heterochomatic proteins are already present in unicellular eukaryotes such as *S. cerevisiae* and *T. thermophila*, while more specific ones are found in mammals, which indeed have more complex repressive chromatin states (Garcia et al. 2007). Similar observations are made in studies focused on the large repertoire of histone modifiers in mammals and in work on PcG proteins. In summary, these studies propose that an elaboration of chromatin states is based on (unique) combinations of histone modifications.

Our phylogenomic profile supports the above idea of regulatory complexification. Indeed, we find that older proteins are more general than recent ones, in the sense that the older proteins tend to be found in multiple types of chromatin. In the eukaryotic cluster (III), proteins linked with histone modification are acetylation/deacetylation proteins (RPD3, DMAP1, SIN3A, PCAF), H3K36me3 reader (MRG15) and H3K4me3 writer (CC10). The multicellular and metazoan clusters (IV and V) highlight complexification of histone modifications throughout eukaryotic evolution: new repressive histone marks appeared, respectively H3K9me3 (SU(VAR)3-9) and H3K27me3 (E(Z)). We confirmed these results through an additional analysis of the conservation of *Drosophila* histone modifiers (Supplementary Figure 4). It is interesting to note that in well-studied unicellular organisms (*T. thermophila*, S. cerevisiae, C. *owczarzaki*), repressive methylated histones H3K9 and H3K27 are often absent or present only at a very low level, while they are abundant in the multicellular fungi *N. crassa* (Garcia et al. 2007; Roudier et al. 2011; Ernst et al. 2011; Jamieson et al. 2013; Sebé-Pedrós et al. 2016). Thus we find diversification of histone marks and the acccompanying proteins, and as mentioned above, that may allow for a more fine-grained regulatory control over the genome.

Connected to the modulation of accessibility through histone modifications, our work also supports new regulatory elements to be linked with the transition to multicellularity. We find that the multicellular and metazoan clusters (IV and V) display the first insulator (DWG, CTCF, CC27) and enhancer binding proteins (JRA). Indeed, enhancers and insulators are mechanistically linked: enhancers being distal regulatory regions, they rely on looping with help of insulators to influence the expression of their targets (Krivega & Dean 2012; Phillips-Cremins & Corces 2013).

Taken together, we affirm the importance of regulatory complexification in the success of multicellular life. Like other studies, our work suggests this regulatory complexification to be linked with the need to control chromatin states and their propagation in an increasingly complex landscape of active and repressive genomic regions.

### Outlook

We have enhanced our understanding of the evolution of the chromatin landscape through the chromatin-associated proteome in *Drosophila*. This is a good starting point, and we need additional studies that focus on different cell types and other species to deepen and broaden that knowledge. For instance, we do not know how chromatin states differ in *Drosophila* over development and between tissues. Tackling other organisms, such as human cell lines, mouse (*M. musculus*), worm (*C. elegans*), and plant (*A. thaliana*), is in principle possible, as the experimental technique that underpins the data set in *Drosophila*, DamID, has been successfully adapted to each of these species (Vogel et al. 2006; Vogel, van Steensel, et al. 2007; Zhang et al. 2007; Gómez-Saldivar et al. 2016; Tosti et al. 2018). Comparing different species is crucial to determine if the evolutionary scenarios that we propose indeed hold true and how they may need to be refined or reconsidered. One future breakthrough we hope for, is that such studies could provide insight into new BLACK-associated proteins and perhaps lead to a better molecular and evolutionary characterization of this type. Moreover, some classes of proteins are better studied in species other than *Drosophila*. For instance, in our dataset, five proteins are responsible for histone acetylation/deacetylation, but substrate specificity and links with previously inferred chromatin states are not well-investigated in fly species. Contrastingly, acetylases (HAT) and deacetylases (HDAC) specificity are well-characterized in human (Seto & Yoshida 2014) and thus *H. sapiens* could be a better subject for questions in this area. Furthermore, non-coding RNAs are tightly associated to both active and inactive chromatin in eukaryotes, including in *S. pombe (Martienssen et al. 2005)*, in various mammals (Saksouk et al. 2015), and in *D. melanogaster (Fagegaltier et al. 2009)*. Thus we advocate for an inclusion of ncRNA functionality within the analyses on different chromatin states across species. Clearly our current study is but an introduction that shows the potential exists for new insights into the evolution of the chromatin landscape.

## Acknowledgements

We thank Joke van Bemmel for help interpreting the genome-wide binding data and the network model of chromatin organization. We thank Guillaume Filion for critical and stimulating comments and additional insight into the data, and Christian Rödelsperger for critical and encouraging comments. We thank Carole Knibbe for statistical advice. AC kindly acknowledges the support by LabEx Memolife and Fondation Bettencourt Schueller.

