## Supplementary information for "Evolution of the *D. melanogaster* chromatin landscape and its associated proteins"

| Species name | Three-character code | Species name (continued) | Code |
|---|---|---|---|
| *Acyrthosiphon pisum* | Api | *Homo sapiens* | Hsa |
| *Aeropyrum pernix* | Ape | *Methanobrevibacter smithii* | Msm |
| *Anabaena variabilis* | Ava | *Methanosarcina acetivorans* | Mac |
| *Anopheles gambiae* | Aga | *Monodelphis domestica* | Mdo |
| *Apis mellifera* | Ame | *Monosiga brevicollis* | Mbr |
| *Arabidopsis thaliana* | Ath | *Mus musculus* | Mmu |
| *Atta cephalotes* | Ace | *Mycoplasma genitalium* | Mge |
| *Bacillus subtilis* | Bsu | *Naegleria gruberi* | Ngr |
| *Bigelowiella natans* | Bna | *Nematostella vectensis* | Nve |
| *Bombyx mori* | Bmo | *Ornithorhynchus anatinus* | Oan |
| *Branchiostoma floridae* | Bfl | *Oryza sativa* | Osa |
| *Buchnera aphidicola* | Bap | *Pediculus humanus* | Phu |
| *Caenorhabditis elegans* | Cel | *Phaeodactylum tricornutum* | Ptr |
| *Candidatus Korarchaeum cryptofilum* | Cko | *Physcomitrella patens* | Ppa |
| *Capitella teleta* | Cte | *Pirellula staleyi* | Pst |
| *Chlamydomonas reinhardtii* | Cre | *Plasmodium falciparum* | Pfa |
| *Ciona intestinalis* | Cin | *Pyrococcus abyssi* | Pab |
| *Cryptococcus neoformans* | Cne | *Saccharomyces cerevisiae* | Sce |
| *Culex quinquefasciatus* | Cqu | *Schistosoma japonicum* | Sja |
| *Danaus plexippus* | Dpl | *Schizosaccharomyces pombe* | Spo |
| *Deinococcus radiodurans* | Dra | *Sulfolobus solfataricus* | Sso |
| *Dictyostelium discoideum* | Ddi | *Tribolium castaneum* | Tca |
| *Drosophila melanogaster* | Dme | *Trichoplax adhaerens* | Tad |
| *Drosophila simulans* | Dsi | | |
| *Emiliania huxleyi* | Ehu | | |
| *Escherichia coli* | Eco | | |
| *Geodermatophilus obscurus* | Gob | | |
| *Glossina morsitans* | Gmo | | |
| *Guillardia theta* | Gth | | |
| *Haloferax volcanii* | Hvo | | |

**Supplementary Table 1.** Species names in alphabetical order and their corresponding three-character code as used in Figure 1 and Supplementary Figure 4.

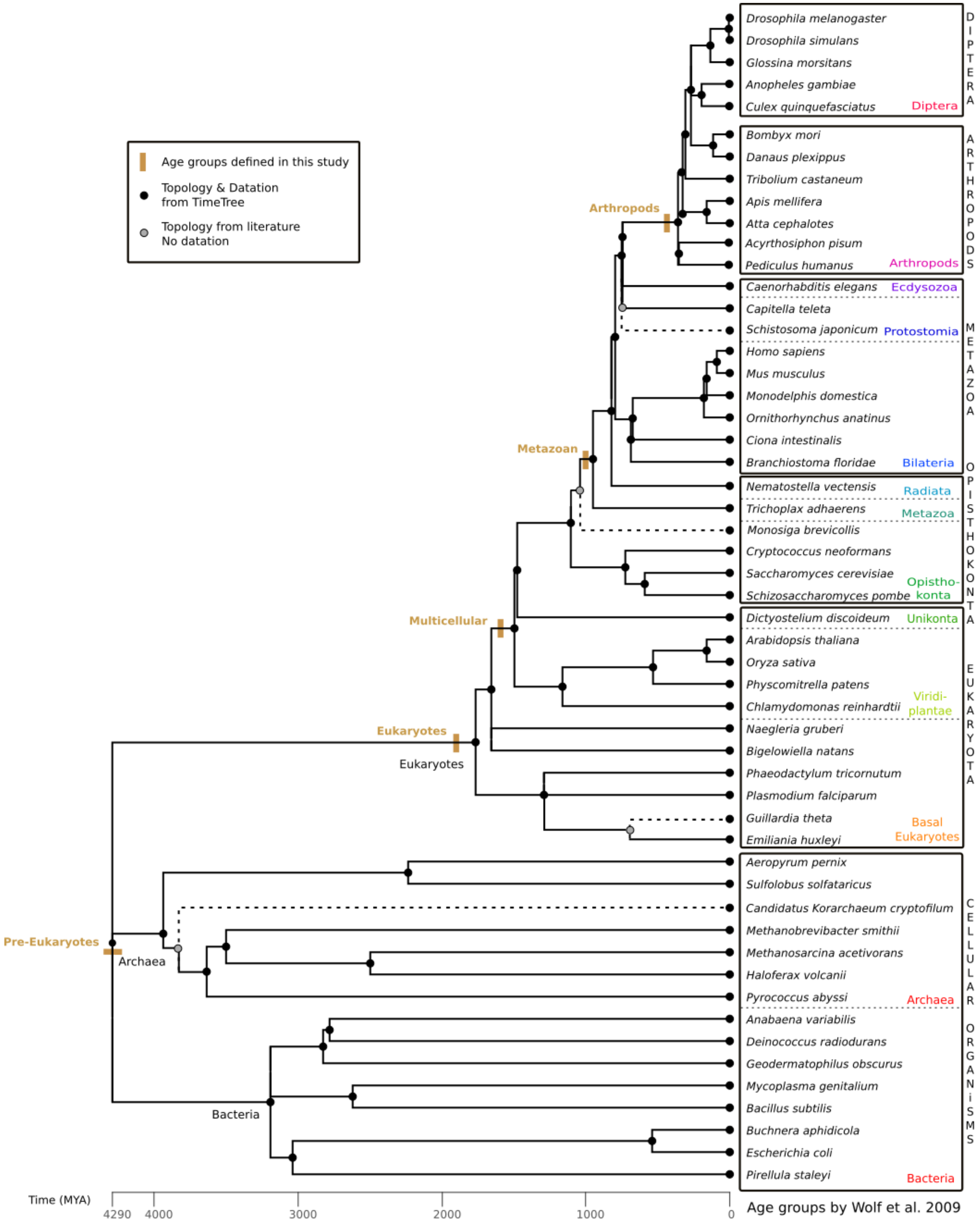

Age groups defined in this study
Topology & Datation from TimeTree
Topology from literature No datation
Drosophila melanogaster
Drosophila simulans
Glossina morsitans
Anopheles gambiae
Culex quinquefasciatus
Diptera
DIPTERA
Bombyx mori
Danaus plexippus
Tribolium castaneum
Apis mellifera
Atta cephalotes
Acyrthosiphon pisum
Pediculus humanus
Arthropods
ARTHROPODS
Caenorhabditis elegans
Ecdysozoa
Capitella teleta
Schistosoma japonicum
Protostomia
Homo sapiens
Mus musculus
Monodelphis domestica
Ornithorhynchus anatinus
Ciona intestinalis
Branchiostoma floridae
Bilateria
METAZOA
Nematostella vectensis
Radiata
Trichoplax adhaerens
Metazoa
Monosiga brevicollis
Cryptococcus neoformans
Saccharomyces cerevisiae
Schizosaccharomyces pombe
Opistho-konta
OPISTHOKONTA
Dictyostelium discoideum
Unikonta
Arabidopsis thaliana
Oryza sativa
Physcomitrella patens
Chlamydomonas reinhardtii
Viridi-plantae
Naegleria gruberi
Bigelowiella natans
Phaeodactylum tricornutum
Plasmodium falciparum
Guillardia theta
Emiliania huxleyi
Basal Eukaryotes
EUKARYOTA
Aeropyrum pernix
Sulfolobus solfataricus
Candidatus Korarchaeum cryptofilum
Methanobrevibacter smithii
Methanosarcina acetivorans
Haloferax volcanii
Pyrococcus abyssi
Archaea
Anabaena variabilis
Deinococcus radiodurans
Geodermatophilus obscurus
Mycoplasma genitalium
Bacillus subtilis
Buchnera aphidicola
Escherichia coli
Pirellula staleyi
Bacteria
CELLULAR ORGANISMS
Arthropods
Metazoan
Multicellular
Eukaryotes
Eukaryotes
Pre-Eukaryotes
Archaea
Bacteria
Time (MYA)
4290
4000
3000
2000
1000
0
Age groups by Wolf et al. 2009

**Supplementary Figure 1.** Phylogenetic species tree of the 53 species as extracted from Timetree (http://www.treetime.org). Age groups defined in this study are represented as brown bars and labels on the tree. As a validation of our groups, on the right we indicate in black boxes the age groups used by (Wolf et al., 2009) to study homologs of *Drosophila melanogaster* proteins. In the boxes, colored names referred to the thirteen species groups that we defined in the main manuscript, see Fig. 1. At the bottom, time is indicated in Millions of Years Ago (MYA), with species divergence times extracted from Timetree. Species absent from TimeTree were manually added based on literature (dashed branches). For those, no dating data were available and nodes are arbitrarily placed on the timescale (grey nodes). Black nodes are dated by Timetree.

| YELLOW | Dollo | | | | |
|---|---|---|---|---|---|
| **PAM** | **Pre-Eukaryotes** | **Eukaryota** | **Opisthokonta** | **Metazoa** | **Arthopods** |
| **Pre-Euk, I+II** | 3 | | | | |
| **Euk, III** | 4 | 11 | | | |
| **Multicell, IV** | 1 | 5 | | | |
| **Metazoa, V** | | 5 | 2 | 3 | |
| **Arthopods, VI** | | 1 | | 2 | 2 |

| RED | Dollo | | | | |
|---|---|---|---|---|---|
| **PAM** | **Pre-Eukaryotes** | **Eukaryota** | **Opisthokonta** | **Metazoa** | **Arthopods** |
| **Pre-Euk, I+II** | 6 | | | | |
| **Euk, III** | 7 | 20 | | | |
| **Multicell, IV** | 3 | 9 | | | |
| **Metazoa, V** | | 11 | 4 | 9 | |
| **Arthopods, VI** | | | 1 | 5 | 8 |

| GREEN | Dollo | | | | |
|---|---|---|---|---|---|
| **PAM** | **Pre-Eukaryotes** | **Eukaryota** | **Opisthokonta** | **Metazoa** | **Arthopods** |
| **Pre-Euk, I+II** | 1 | | | | |
| **Euk, III** | 1 | 3 | | | |
| **Multicell, IV** | | 2 | | | |
| **Metazoa, V** | | 6 | 1 | | |
| **Arthopods, VI** | | 1 | 1 | 2 | 7 |

**Supplementary Table 2.** Overlap between clustering by Partitioning Around Medoids (PAM) and the Dolly parsimony method for three chromatin types, respectively YELLOW, RED, and GREEN. Numbers indicate the number of shared chromatin-associated proteins, empty cells signal zero shared proteins. See also Figure 3 and 5B in the main text, and Supplementary Table 3.

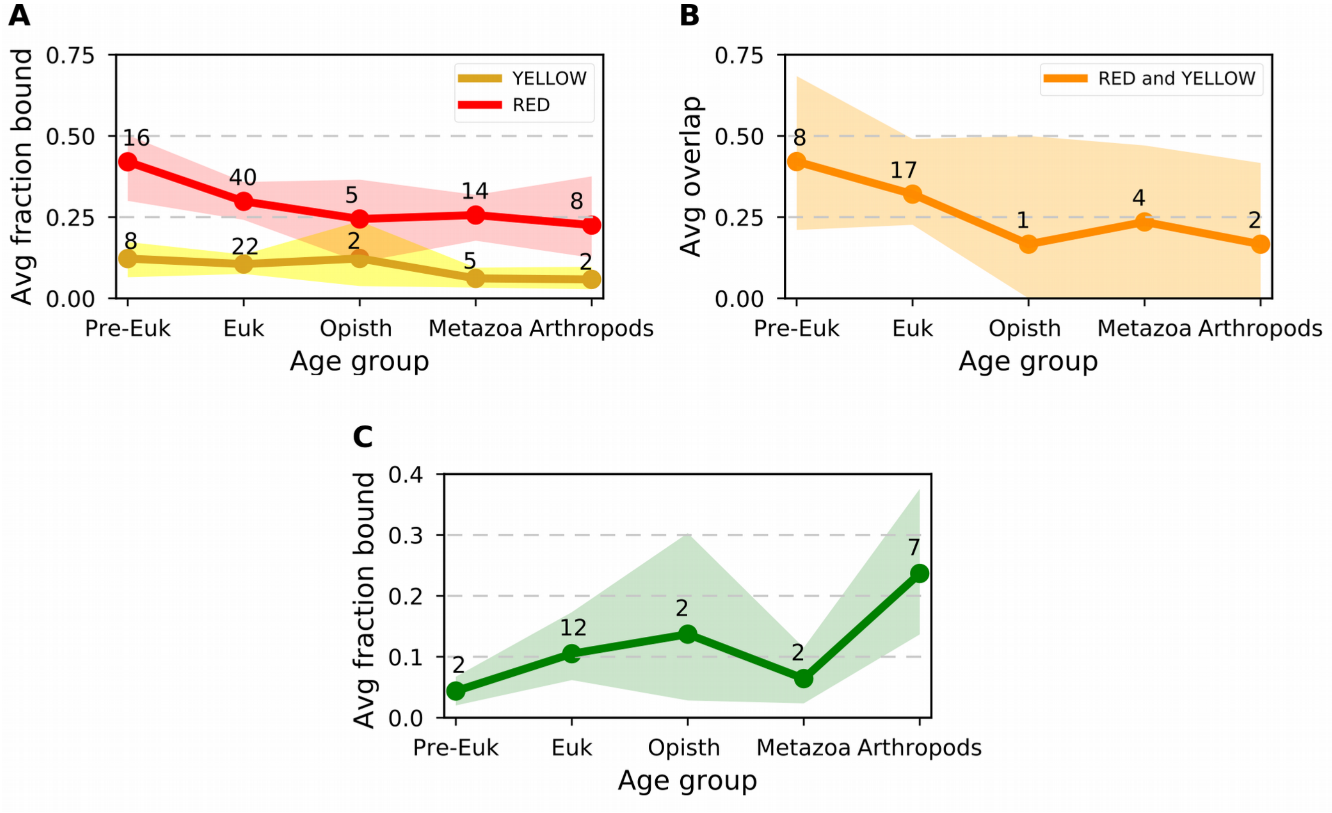


**Supplementary Figure 2.** Average fraction of genome bound by proteins over evolutionary age groups. The evolutionary age is determined by Dollo parsimony. In contrast in Fig. 3, we use Partitioning Around Medoids. **(A)** RED and YELLOW fraction bound. **(B)** Average overlap between RED and YELLOW. **(C)** GREEN fraction bound. See Methods and Fig. 3 in the main text for details.

| DB[1] | A[2] | N[3] | O[4] | U[5] | AVGi[6] | AVGb[7] | Mann-Whitney Test |
|---|---|---|---|---|---|---|---|
| Panther7 | W | 38 | E* | - | 822.0 | 670.3 | U = 6.6e+05 (p = 0.0741) |
| Multiparanoid | W | 12 | E***/O* | Dm* | 854.8 | 781.7 | U = 6.3e+05 (p = 0.0161) |
| Jaccard | W | 12 | E*** | Dm* | 1145.3 | 1015.9 | U = 6.3e+05 (p = 0.0109) |
| Lens | W | 12 | E*** | C*/Dm* | 1145.3 | 1015.9 | U = 6.3e+05 (p = 0.0109) |
| OthoMCL | W | 12 | E*/O* | - | 630.9 | 622.7 | U = 6.7e+05 (p = 0.129) |
| Panther7 | D | 38 | E* | Di** | 1329.4 | 1114.2 | U = 6.2e+05 (p = 0.00757) |
| Multiparanoid | D | 12 | E*** | C*/Dm** | 1018.4 | 937.5 | U = 6.3e+05 (p = 0.0111) |
| Jaccard | D | 12 | E*** | Dm*** | 1316.7 | 1154.8 | U = 6.1e+05 (p = 0.00341) |
| Lens | D | 12 | E*** | C*/Dm*** | 1155.6 | 1076.8 | U = 6.2e+05 (p = 0.00596) |
| OthoMCL | D | 12 | E*** | C*/Dm* | 880.7 | 817.9 | U = 6.4e+05 (p = 0.0191) |

**Supplementary Table 3.** Age enrichment tests for *D. melanogaster* CAPs using different algorithms. The column headings indicate the following, from left to right. DB: The database is DROME_PPODv4, clustered with the corresponding method. A: The algorithm used for enrichment tests, Wagner (W) and Dollo (D). N: Number of species in the species tree. O: overrepresented age groups, Eukaryota (E) and Opisthokonta (O). U: under-represented age groups, *D. melanogaster* (Dm), Cellular organism (C), and Diptera (Di). AVGi: average age input set. AVGb: average age background set. Fisher's exact test was used to calculate the significance of the differences for each age group: *P < 0.05; **P < 0.01; ***P < 0.001.

| Cluster name | Number | GO Identifier | GO Description | FDR | log10(FDR) |
|---|---|---|---|---|---|
| Chromatin organization | 2 | GO:0051276 | chromosome organization | 1.000E-12 | -12.0000 |
| | 16 | GO:0006325 | chromatin organization | 1.000E-12 | -12.0000 |
| | 27 | GO:0071824 | protein-DNA complex subunit organization | 1.374E-07 | -6.8620 |
| Regulation of cell cycle | 14 | GO:0030522 | intracellular receptor signaling pathway | 4.149E-02 | -1.3821 |
| | 25 | GO:0051130 | positive regulation of cellular component organization | 7.371E-03 | -2.1325 |
| | 26 | GO:0048585 | negative regulation of response to stimulus | 1.101E-03 | -2.9583 |
| | 29 | GO:0051129 | negative regulation of cellular component organization | 3.728E-02 | -1.4285 |
| | 31 | GO:0051726 | regulation of cell cycle | 6.825E-04 | -3.1659 |
| Regulation of transcription | 1 | GO:0051172 | negative regulation of nitrogen compound metabolic process | 1.000E-12 | -12.0000 |
| | 19 | GO:0040029 | regulation of gene expression, epigenetic | 1.847E-11 | -10.7336 |
| | 23 | GO:0010628 | positive regulation of gene expression | 5.013E-07 | -6.2999 |
| | 33 | GO:0009890 | negative regulation of biosynthetic process | 1.000E-12 | -12.0000 |
| | 34 | GO:0051173 | positive regulation of nitrogen compound metabolic process | 1.433E-06 | -5.8439 |
| | 35 | GO:0009891 | positive regulation of biosynthetic process | 1.433E-06 | -5.8439 |
| | 36 | GO:0010629 | negative regulation of gene expression | 1.000E-12 | -12.0000 |
| Transcription | 15 | GO:0006259 | DNA metabolic process | 9.703E-09 | -8.0131 |
| | 17 | GO:0006352 | DNA-templated transcription, initiation | 6.473E-03 | -2.1889 |
| | 24 | GO:0031399 | regulation of protein modification process | 2.428E-03 | -2.6147 |
| Protein modification | 8 | GO:0018193 | peptidyl-amino acid modification | 1.290E-05 | -4.8894 |
| | 18 | GO:0098732 | macromolecule deacylation | 1.556E-02 | -1.8079 |
| | 21 | GO:0008213 | protein alkylation | 1.864E-03 | -2.7295 |
| | 22 | GO:0043543 | protein acylation | 3.695E-05 | -4.4324 |
| Response to stress | 3 | GO:0006974 | cellular response to DNA damage stimulus | 6.589E-05 | -4.1812 |
| | 11 | GO:0033993 | response to lipid | 6.825E-04 | -3.1659 |
| | 28 | GO:0014070 | response to organic cyclic compound | 4.149E-02 | -1.3821 |
| other | 0 | GO:0002376 | immune system process | 1.885E-03 | -2.7246 |
| | 6 | GO:0008283 | cell proliferation | 4.531E-02 | -1.3438 |
| | 7 | GO:0032259 | methylation | 1.910E-03 | -2.7190 |
| | 9 | GO:0007059 | chromosome segregation | 3.728E-02 | -1.4285 |
| Development | 5 | GO:0061061 | muscle structure development | 1.024E-02 | -1.9897 |

| | | | | | |
|---|---|---|---|---|---|
| | 10 | GO:0002164 | larval development | 3.762E-02 | -1.4246 |
| | 13 | GO:0007389 | pattern specification process | 2.671E-02 | -1.5734 |
| | 20 | GO:0007417 | central nervous system development | 4.149E-02 | -1.3821 |
| Cell cycle | 4 | GO:1903047 | mitotic cell cycle process | 3.164E-05 | -4.4997 |
| | 12 | GO:0016358 | dendrite development | 1.433E-06 | -5.8439 |
| | 30 | GO:0033301 | cell cycle comprising mitosis without cytokinesis | 2.414E-02 | -1.6172 |
| | 32 | GO:0048667 | cell morphogenesis involved in neuron differentiation | 9.506E-04 | -3.0220 |

**Supplementary Table 4.** Gene Ontology enrichment analysis of CAPs. We used WebGestalt to search 107 chromatin-associating proteins against a *D. melanogaster* background for enriched terms. The resulting 37 GO terms were mapped onto a two-dimensional semantic space and clustered into 9 groups. These groups have been manually named in the first column. We labelled each GO term with a number for visualization purposes in Figure 2A, see the second column. The third and fourth column give the GO term identifier and its accompanying short description. False Discovery Rates (FDR), and their log10 values are reported in column five and six, and are used to colour the circles in Figure 2A. Any GO terms with log10(FDR) < -12 were set to -12. See the main manuscript for details (Methods and Results).

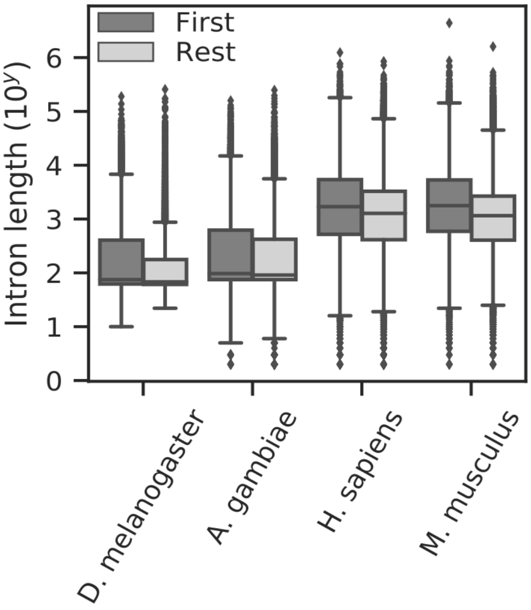


**Supplementary Figure 3.** Intron length in four species, the fruit fly *D. melanogaster*, the mosquito *A. gambiae*, human *H. sapiens*, and mouse *M. musculus*. For each gene, we take the first two 5' introns as "First" and the following introns as "Rest". In all species, the first introns are significantly longer than the rest introns. Note the logarithmic scale on the y-axis.

| | *D. melanogaster* | *A. gambiae* | *H. sapiens* | *M. musculus* |
|---|---|---|---|---|
| **A. First vs rest** | | | | |
| Cellular Organism | **0.000134** | **0.000892** | 0.0238 | **<1e-5** |
| Eukaryota | **<1e-5** | **<1e-5** | **<1e-5** | **<1e-5** |
| Opisthokonta | **0.000154** | 0.044 | **<1e-5** | **<1e-5** |
| Bilateria | **<1e-5** | **3.93e-5** | **<1e-5** | **<1e-5** |
| Diptera / Mammalia | **<1e-5** | **0.000106** | **<1e-5** | **<1e-5** |
| Species-specific | **0.000121** | 0.581 | **0.00205** | 0.347 |
| **B. First YELLOW vs RED** | | | | |
| Cellular Organism | 0.0233 | 0.0489 | 0.966 | 0.114 |
| Eukaryota | **<1e-5** | **0.00212** | 0.153 | 0.0231 |
| Opisthokonta | **0.00033** | 0.108 | 0.108 | **0.00893** |
| Bilateria | **<1e-5** | **3.25e-5** | 0.153 | **0.00452** |
| Diptera / Mammalia | **<1e-5** | **0.00033** | 0.287 | 0.0364 |
| Species-specific | **<1e-5** | **0.000259** | 0.404 | 0.772 |
| **C. Intron ratio YELLOW vs RED** | | | | |
| Cellular Organism | 0.0406 | 0.0474 | 0.602 | 0.0166 |
| Eukaryota | **0.000286** | 0.038 | 0.344 | 0.123 |
| Opisthokonta | 0.123 | 0.123 | 0.587 | 0.127 |
| Bilateria | **<1e-5** | 0.0166 | 0.173 | 0.0857 |
| Diptera / Mammalia | **<1e-5** | 0.123 | 0.224 | 0.0572 |
| Species-specific | **<1e-5** | 0.112 | 0.858 | 0.843 |

**Supplementary Table 5.** Three sets of statistical tests to establish significant differences in length (i.e. number of nucleotides) between sets of introns across evolutionary age groups. We show FDR-corrected p-values (alpha=0.01) of Mann-Whitney-U tests, with significant p-values in bold font on a light-grey background. **A.** For all genes, the length of their first two 5' introns is compared to the length of downstream (3') introns. The former are named "first", the latter "rest". Genes are categorized by evolutionary age as estimated by ProteinHistorian. Significant p-values indicate that first introns are longer than rest introns. See main text for details. **B.** For genes in YELLOW and RED, we take the first two 5' introns and compare their length. Again, genes are categorized by evolutionary age. Here significant p-values indicate RED first introns are longer than YELLOW ones. **C.** For genes in YELLOW and RED, we take the ratio of first to rest introns. We then compare ratios and a significant p-value indicates a higher intron length ratio for RED. This means that RED genes have longer first introns relative to their subsequent introns than YELLOW genes do.

**A**

| Chromatin type | Interesting genes | | Reference genes | | Significant GO terms |
|---|---|---|---|---|---|
| YELLOW | $log_2$ ratio ~0.0 | n=697 | all YELLOW | n=1286 | Ribonucleoprotein complex biogenesis; ncRNA metabolic process; DNA metabolic process; cellular response to DNA damage stimulus |
| | $log_2$ ratio >1.0 | n=589 | | | Localization of cell; taxis; appendage development; wing disc development; post-embryonic animal organ morphogenesis; actin filament-based process; embryonic morphogenesis |
| RED | $log_2$ ratio ~0.0 | n=85 | all RED | n=339 | ns |
| | $log_2$ ratio >1.0 | n=254 | | | ns |

**B**

| Intron ratio | Interesting genes | | Reference genes | | Significant GO terms |
|---|---|---|---|---|---|
| $log_2$ ratio ~0.0 | YELLOW | n=697 | all $log_2$ ratio ~0.0 | n=782 | ns |
| | RED | n=85 | | | ns |
| $log_2$ ratio >1.0 | YELLOW | n=589 | all $log_2$ ratio >1.0 | n=843 | ns |
| | RED | n=254 | | | Negative regulation of developmental process; negative regulation of multicellular organismal process |

**C**

| | Interesting genes | | Reference genes | | Significant GO terms |
|---|---|---|---|---|---|
| Chromatin | all YELLOW | n=1286 | all | n=1625 | mRNA metabolic process; DNA metabolic process; RNA splicing; cellular response to DNA damage stimulus |
| | all RED | n=339 | | | Negative regulation of developmental process; negative regulation of multicellular organismal process; post-embryonic animal organ morphogenesis |
| Intron ratio | all $log_2$ ratio ~0.0 | n=782 | | | Ribonucleoprotein complex biogenesis; RNA localization; ncRNA metabolic process; DNA metabolic process; nucleic acid phosphodiester bond hydrolysis; cellular response to DNA damage stimulus |
| | all $log_2$ ratio >1.0 | n=843 | | | post-embryonic animal organ morphogenesis; appendage development; wing disc development; muscle structure development; taxis; regulation of nervous system development; response to growth factor; embryonic morphogenesis; localization of cell; actin filament-based process; growth; cell morphogenesis involved in neuron differentiation; circulatory system development; exocrine system development; |

**Supplementary Table 6.** Relative GO enrichment analysis for YELLOW and RED genes using WebGestalt. GO Terms are reported if they have an FDR < 0.004, which is a reasonable limit considering the number of tests (n=12). ns = no significant term.

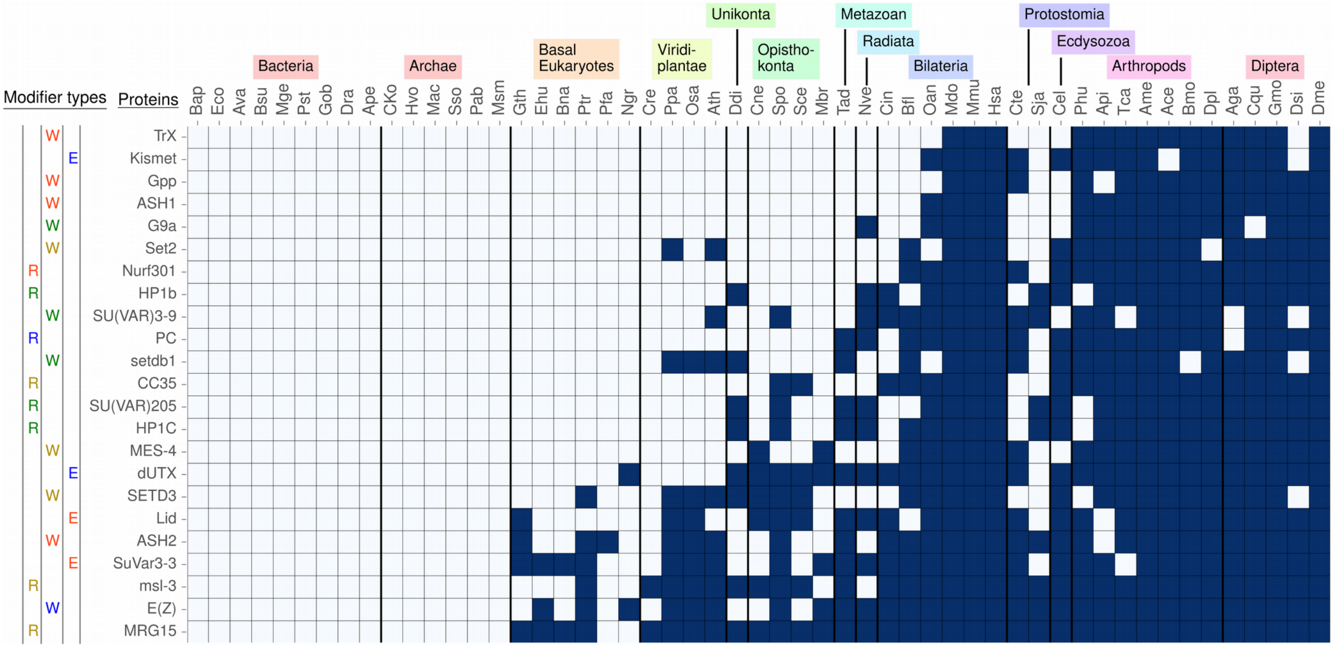


**Supplementary Figure 4.** Phylogenetic profile of *D. melanogaster* histone mark modifiers. On the left, reader (R), writer (W), and eraser (E) proteins are specified with a color code corresponding to the chromatin types and their typical histone modifications. YELLOW refers to H3K36me3, YELLOW and RED are shown in orange and indicate H3K4me3, BLUE is H3K27me3, GREEN is H3K9me2/3. In the matrix, dark blue rectangles represent the presence of a homolog, grey rectangles its absence. On top, 13 species groups are defined to aid the reader and the three letter codes refer to species names as given in Supplementary Table 1.

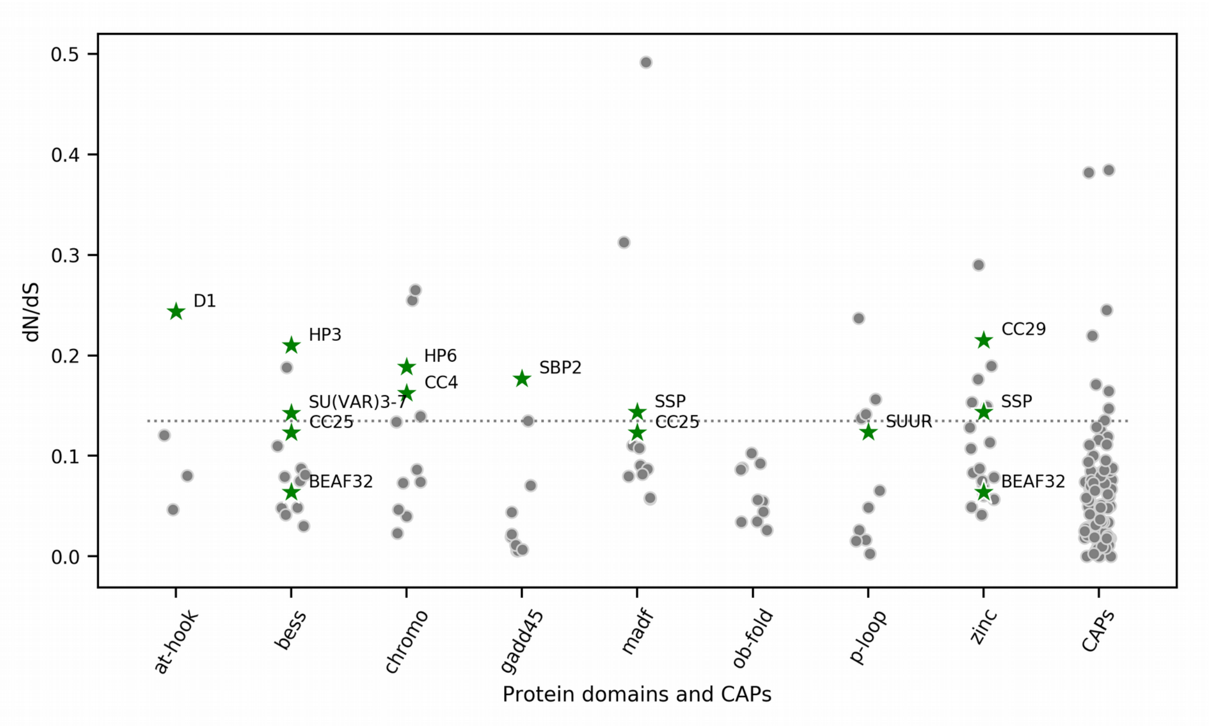


**Supplementary Figure 5.** dN/dS values for randomly selected proteins with the same domains as young GREEN CAPs. As a reference the dN/dS of all chromatin-associated proteins (CAPs) are shown on the right (see also Figure 5C). Using InterPro we extracted the following domains for young GREEN proteins. D1: AT-hook, BEAF32: Zinc finger and BESS, HP3 (LHR): BESS, SU(VAR)3-7: BESS, HP6: Chromo, SUUR: P-loop, SSP: Zinc finger and MADF, CC4: Chromo, CC29: Zinc finger, SBP2: Gadd45, CC25: BESS and MADF. dN/dS values were calculated using the same set of *drosophilids* as for the CAPs, see Methods and Results in the main manuscript for details.

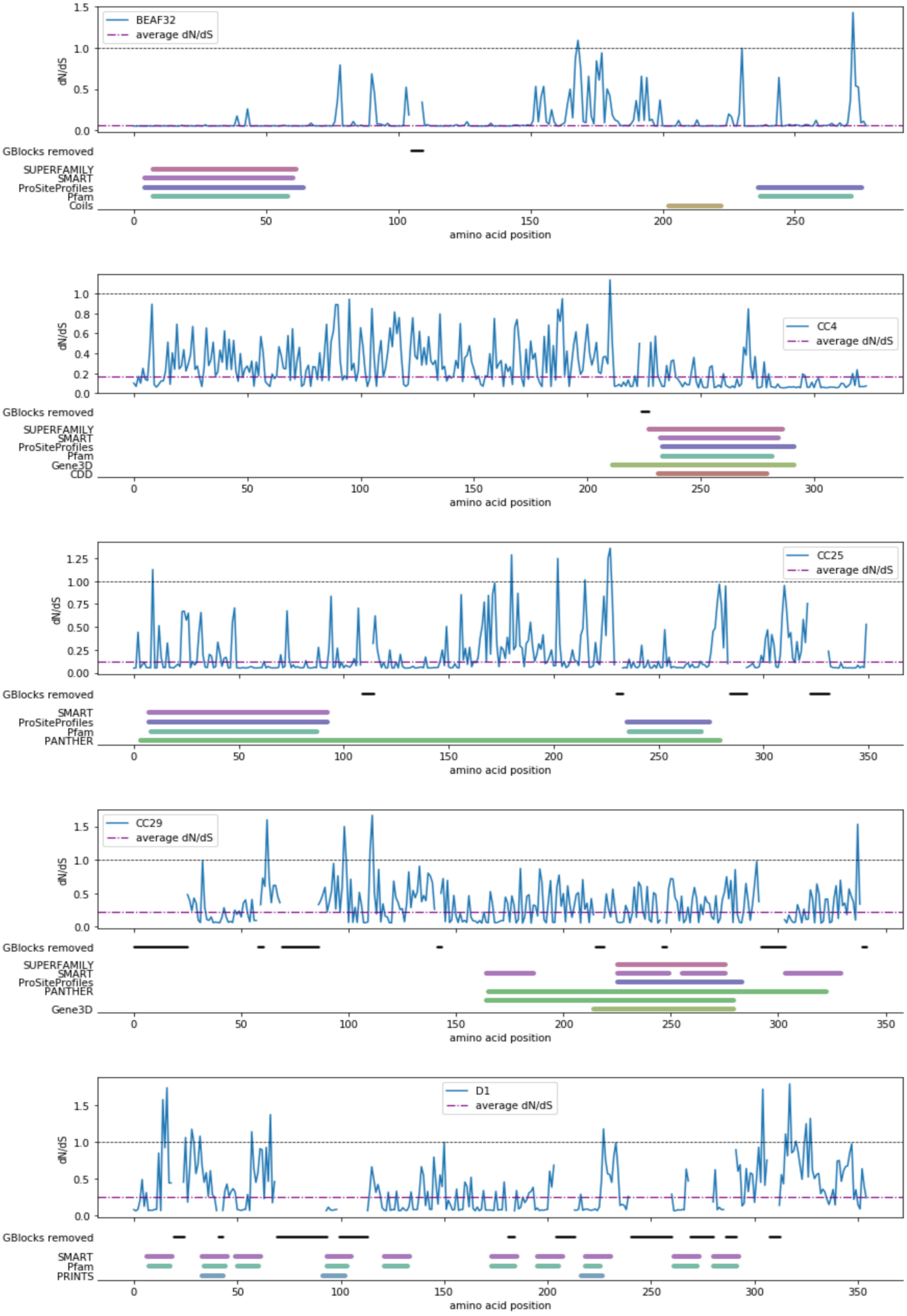
BEAF32
average dN/dS
dN/dS
GBlocks removed
SUPERFAMILY
SMART
ProSiteProfiles
Pfam
Coils
amino acid position
CC4
average dN/dS
dN/dS
GBlocks removed
SUPERFAMILY
SMART
ProSiteProfiles
Pfam
Gene3D
CDD
amino acid position
CC25
average dN/dS
dN/dS
GBlocks removed
SMART
ProSiteProfiles
Pfam
PANTHER
amino acid position
CC29
average dN/dS
dN/dS
GBlocks removed
SUPERFAMILY
SMART
ProSiteProfiles
PANTHER
Gene3D
amino acid position
D1
average dN/dS
dN/dS
GBlocks removed
SMART
Pfam
PRINTS
amino acid position

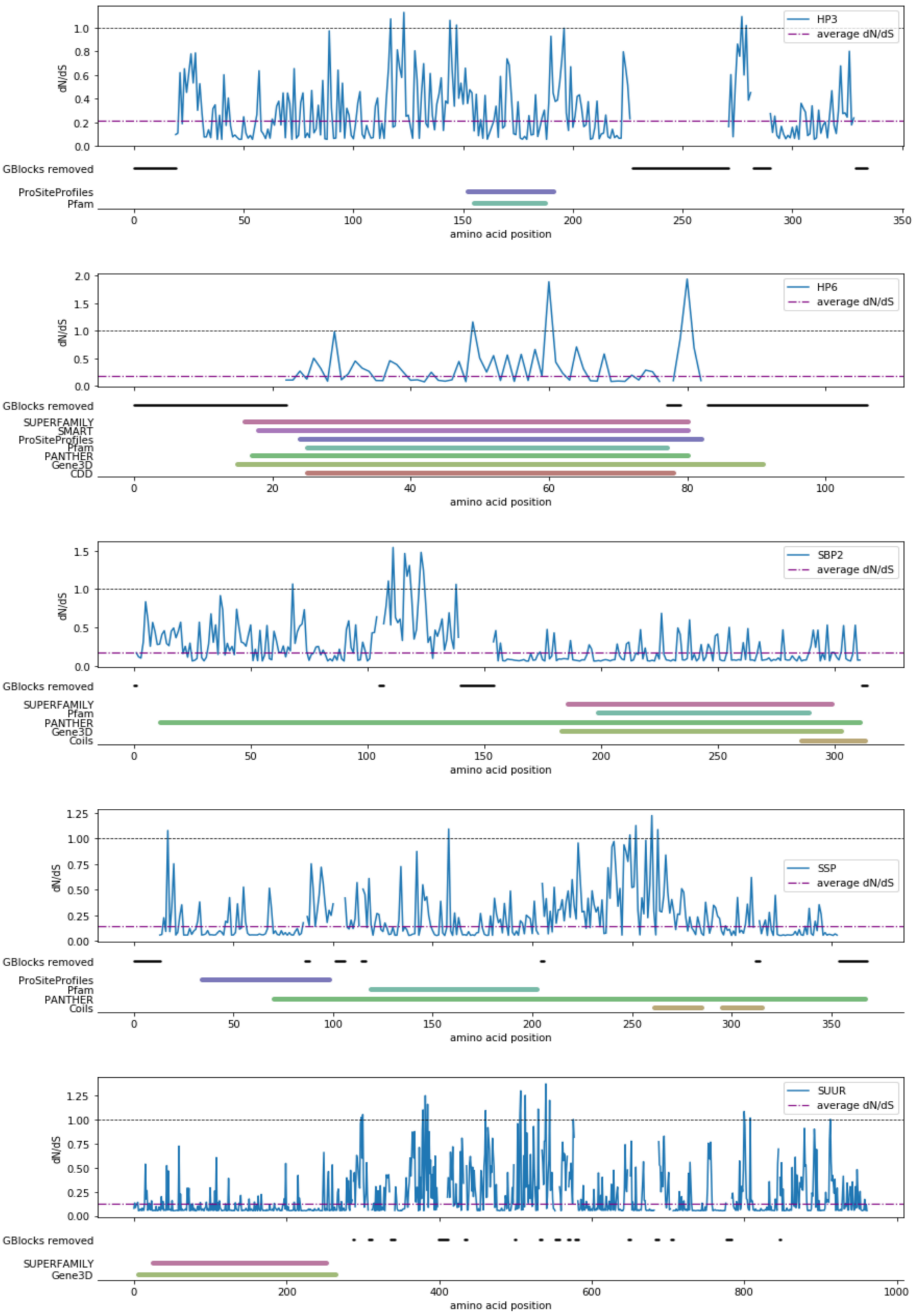
HP3
average dN/dS
dN/dS
GBlocks removed
ProSiteProfiles
Pfam
amino acid position
HP6
average dN/dS
dN/dS
GBlocks removed
SUPERFAMILY
SMART
ProSiteProfiles
Pfam
PANTHER
Gene3D
CDD
amino acid position
SBP2
average dN/dS
dN/dS
GBlocks removed
SUPERFAMILY
Pfam
PANTHER
Gene3D
Coils
amino acid position
SSP
average dN/dS
dN/dS
GBlocks removed
ProSiteProfiles
Pfam
PANTHER
Coils
amino acid position
SUUR
average dN/dS
dN/dS
GBlocks removed
SUPERFAMILY
Gene3D
amino acid position

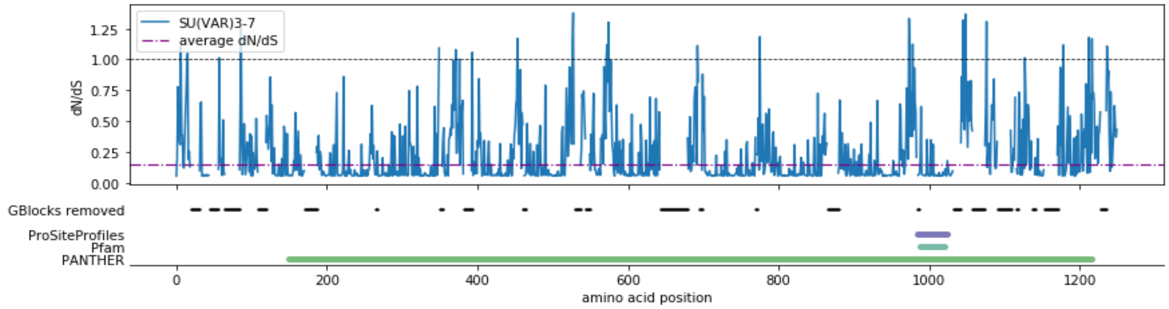


**Supplementary Figure 6.** Summary of dN/dS and Interpro predicted protein domains along the protein's amino acid sequence for the 11 GREEN arthropod proteins. In each panel, the protein's average dN/dS is given as a purple horizontal line and amino acid dN/dS scores by the blue line. Below each panel, the amino acids removed by Gblocks are indicated by black horizontal lines, followed by Interpro protein domain predictions, that originate from a range of databases and prediction tools mentioned as labels on the left. Some proteins (notably CC4, SBP2, and SUUR) show low dN/dS for predicted domains and high dN/dS for unstructured parts of the sequence. Yet, others do not show such a pattern, for instance CC29, D1, HP3 and HP6. Note that HP3 is also known as LHR.

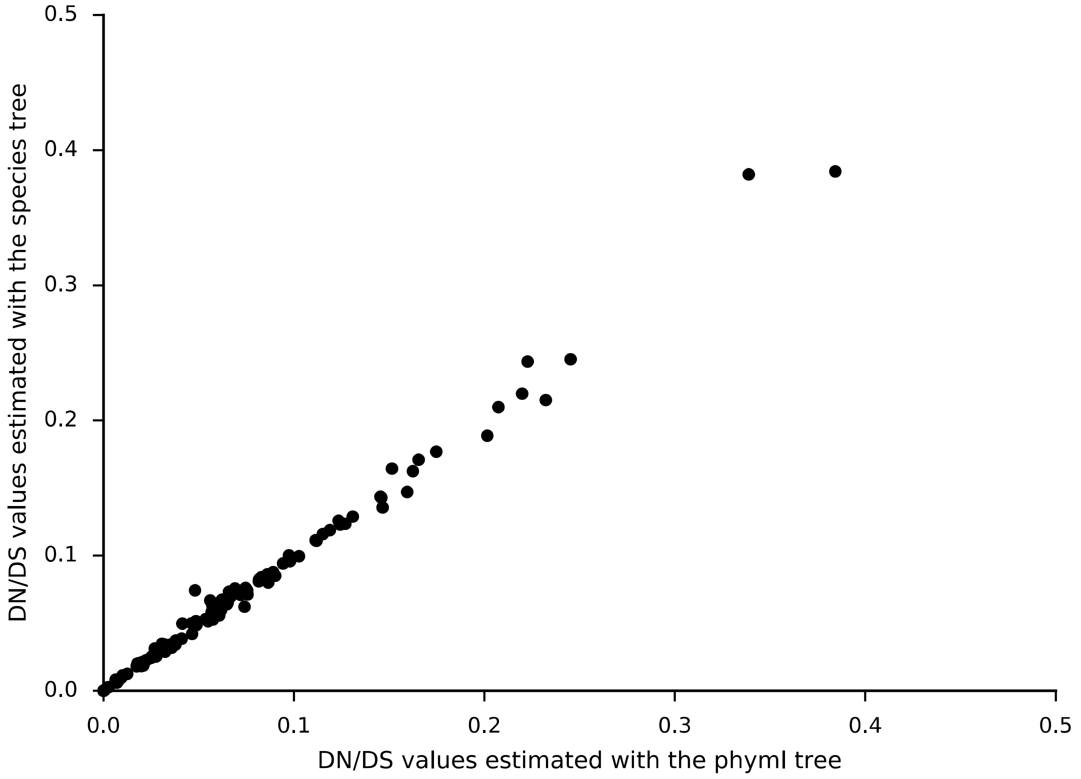


**Supplementary Figure 7.** Scatter plot of chromatin protein dN/dS ratios obtained using a gene tree topology (PhyML) and the Species tree topology (Timetree).

| Foreground branch | Likelihood | | 2DL | Degrees of freedom | p-value |
|---|---|---|---|---|---|
| | Null Model | Model A | | | |
| **1** | **-5180.81** | **-5174.19** | **13.24** | **1** | **$4.91 . 10^{-3}$** |
| 2 | -5181.25 | -5181.63 | -0.76 | 1 | 1 |
| 3 | -5178.25 | -5175.12 | 6.26 | 1 | 0.22 |
| **4** | **-5180.81** | **-5174.19** | **13.24** | **1** | **$4.91 . 10^{-3}$** |
| **5** | **-5180.21** | **-5167.94** | **24.54** | **1** | **$1.31 . 10^{-5}$** |
| 6 | -5181.38 | -5177.30 | 8.16 | 1 | 0.07 |
| **7** | **-5180.17** | **-5172.50** | **15.34** | **1** | **$1.62 . 10^{-3}$** |
| 8 | -5181.47 | -5181.63 | -0.32 | 1 | 1 |
| 9 | -5181.63 | -5181.63 | 0.00 | 1 | 1 |
| **10** | **-5180.24** | **-5162.70** | **35.08** | **1** | **$5.68 . 10^{-8}$** |
| 11 | -5176.93 | -5174.40 | 5.06 | 1 | 0.44 |
| 12 | -5178.42 | -5173.55 | 9.74 | 1 | 0.032 |
| 13 | -5181.63 | -5181.63 | 0.00 | 1 | 1 |
| 14 | -5181.63 | -5181.63 | 0.00 | 1 | 1 |
| 15 | -5181.41 | -5181.60 | -0.38 | 1 | 1 |
| 16 | -5181.63 | -5181.63 | 0.00 | 1 | 1 |
| 17 | -5180.50 | -5181.47 | -1.94 | 1 | 1 |
| 18 | -5181.63 | -5181.63 | 0.00 | 1 | 1 |

**Supplementary Table 7.** Summary of positive selection tests under the free-ratio model and branch-site model. Foreground branch numbers refer to a top-to-bottom and left-to-right numbering of the branches of the tree in Figure 6A. The bold font on a grey background indicates significant selection tests. P-values are adjusted for multiple testing with the Bonferroni correction.

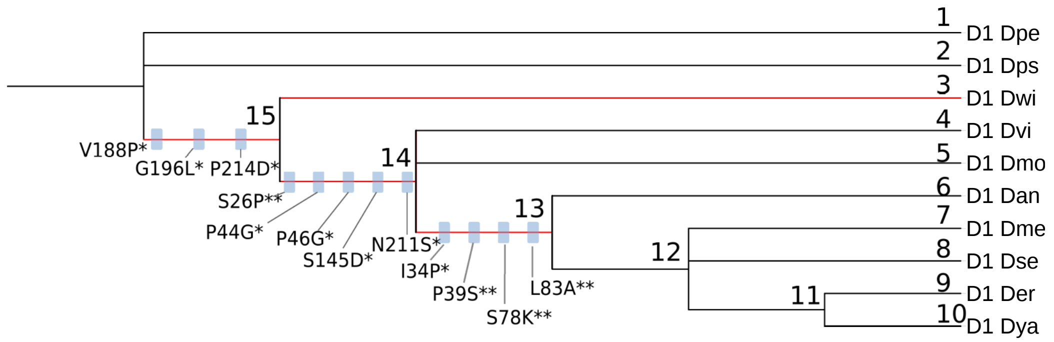


**Supplementary Figure 8.** Gene tree of D1 constructed with Phyml (with arbitrary branch lengths). The four branches with positive selection events are highlighted in red (p< 0.01, Bonferroni correction). On branches with more than one positively selected site, blue boxes indicate the amino acid substitution under positive selection, with the significance given as posterior probability of dN/dS > 1 (* for Pr > 0.95, ** for Pr > 0.99). Species are indicated by 3 letter abbreviations, see Figure 6A in the main text.

| Foreground branch | Likelihood | | 2DL | Degrees of freedom | p-value |
|---|---|---|---|---|---|
| | Null Model | Model A | | | |
| 1 | -5198.60 | -5198.60 | 0.00 | 1 | 1 |
| 2 | -5198.60 | -5198.60 | 0.00 | 1 | 1 |
| **3** | **-5195.26** | **-5188.23** | **14.06** | **1** | **$2.65 . 10^{-3}$** |
| 4 | -5194.61 | -5192.83 | 3.56 | 1 | 0.88 |
| 5 | -5198.53 | -5198.04 | 0.98 | 1 | 1 |
| 6 | -5192.72 | -5190.66 | 4.12 | 1 | 0.63 |
| 7 | -5198.49 | -5198.49 | 0.00 | 1 | 1 |
| 8 | -5198.60 | -5198.60 | 0.00 | 1 | 1 |
| 9 | -5198.12 | -5198.12 | 0.00 | 1 | 1 |
| 10 | -5198.60 | -5181.63 | 0.00 | 1 | 1 |
| 11 | -5198.60 | -5198.60 | 0.00 | 1 | 1 |
| 12 | -5196.71 | -5194.95 | 3.52 | 1 | 0.91 |
| **13** | **-5194.00** | **-5184.47** | **19.06** | **1** | **$1.91 . 10^{-4}$** |
| **14** | **-5193.77** | **-5187.46** | **12.62** | **1** | **$5.74 . 10^{-3}$** |
| **15** | **-5195.54** | **-5188.97** | **13.14** | **1** | **$4.36 . 10^{-3}$** |

**Supplementary Table 8.** Summary of positive selection tests using the gene tree topology of Supplementary Figure 4. Foreground branch numbers refer to branches of the tree. The bold font on a grey background indicates significant selection tests. P-values are adjusted for multiple testing with the Bonferroni correction.
